# Masked features of task states found in individual brain networks

**DOI:** 10.1101/2021.06.12.448198

**Authors:** Alexis Porter, Ashley Nielsen, Megan Dorn, Ally Dworetsky, Donnisa Edmonds, Caterina Gratton

**Author notes:** **Correspondence:** Caterina Gratton. **Author Contributions:** A.P., conceptualization, investigation, software, formal analysis, visualization, writing (original draft, review, and editing). A.N., conceptualization, methodology, writing (review and editing). M.D., data collection, task creation. A.D., software, formal analysis. D.E., preliminary analysis of secondary dataset. C.G., conceptualization, methodology, investigation, writing (original draft, review, and editing), resources, supervision. **Competing interests:** The authors have no competing interests.

## Abstract

Completing complex tasks requires that we flexibly integrate information across brain areas. While studies have shown how functional networks are altered during different tasks, this work has generally focused on a cross-subject approach, emphasizing features that are common across people. Here we used extended sampling “precision” fMRI data to test the extent to which task states generalize across people or are individually-specific. We trained classifiers to decode state using functional network data in single-person datasets across 5 diverse task states. Classifiers were then tested on either independent data from the same person or new individuals. Individualized classifiers were able to generalize to new participants. However, classification performance was significantly higher within a person, a pattern consistent across model types, people, tasks, feature subsets, and even for decoding very similar task conditions. Notably, these findings also replicated in a new independent dataset. These results suggest that individual-focused approaches can uncover robust features of brain states, including features obscured in cross-subject analyses. Individual-focused approaches have the potential to deepen our understanding of brain interactions during complex cognition.

**Citation Diversity Statement:** Recently, the field of neuroscience has reported a bias in citation practices such that papers from minority groups are more often under-cited relative to the number of papers in the field (Dworkin et al. 2020). The authors of this paper were proactive in consideration of selecting references that reflect diversity of the field in thought, contribution, and gender. Utilizing previously derived databases (Dworkin et al. 2020; Zhou et al. 2020) we obtained the predicted gender of authors referenced in this manuscript. By this measure (and excluding self-citations to the first and last authors of our current paper), our references contain 13.87% woman(first)/woman(last), 23.3% man/woman, 23.3% woman/man, and 39.53% man/man. This method is limited in that a) names, pronouns, and social media profiles used to construct the databases may not, in every case, be indicative of gender identity and b) it cannot account for intersex, non-binary, or transgender people. Second, we obtained the predicted racial/ethnic category of the first and last author of each reference by databases that store the probability of a first and last name being carried by an author of color(Ambekar et al. 2009). By this measure (and excluding self-citations), our references contain 10.83% author of color (first)/author of color(last), 10.64% white author/author of color, 23.55% author of color/white author, and 54.98% white author/white author. This method is limited in that a) names and Florida Voter Data to make the predictions may not be indicative of racial/ethnic identity, and b) it cannot account for Indigenous and mixed-race authors, or those who may face differential biases due to the ambiguous racialization or ethnicization of their names. We look forward to future work that could help us to better understand how to support equitable practices in science.

## Introduction

Achieving task goals requires the integration of functions associated with many different brain regions organized into large-scale brain networks (Cole et al. 2014; Bertolero et al. 2015 Nov 20). In healthy humans, large-scale network interactions can be measured across different states using functional connectivity MRI (FC), by correlating activity of brain regions over time. Most past studies have focused on measuring network changes during tasks by grouping data across participants, focusing on common task state features (Betti et al. 2013; Cole et al. 2014; Gratton et al. 2016; Rosenberg et al. 2018). These studies demonstrate that relative to rest, functional networks are altered subtly but significantly during different task states (Cole et al. 2013; Krienen et al. 2014; Gratton et al. 2016; Gratton et al. 2018). Machine learning can be used on these subtle network signals to classify task states with good accuracy (Shirer et al. 2012; Cole et al. 2013; Alnæs et al. 2015; Gonzalez-Castillo et al. 2015; Greene et al. 2018; Greene et al. 2020; Rosenberg et al. 2020; Wu et al. 2020), even during self-driven tasks (Shirer et al. 2012; Krienen et al. 2014). However, the logic in many of these past studies implicitly assumes that functional networks are altered in similar ways across individuals during tasks: only network features that are similar in anatomical localization and form will be readily apparent in cross-subject data and of use in predicting task state in new individuals. Here we asked to what extent brain network features during tasks generalize across individuals relative to showing individual-specific characteristics.

One approach to measuring individualized brain function is to shift the focus of data collection and analysis to single individuals through extended single-subject data collection, an approach termed precision fMRI (Gordon, Laumann, Gilmore, et al. 2017; Gratton et al. 2020). This method produces reliable network maps that are sensitive to individual features (Braga and Buckner 2017; Gordon, Laumann, Adeyemo, Gilmore, Nelson, Nico U.F. Dosenbach, and Petersen 2017; Gratton et al. 2018; Braga et al. 2019; Seitzman et al. 2019; DiNicola et al. 2020). We recently used precision fMRI data to demonstrate that brain networks are largely stable within an individual and subject to only subtle modulations during task states (Gratton et al. 2018). These task-modulations appeared to vary across people (Gratton et al. 2018), consistent with reports that task and individual differences variables interact in how they relate to brain networks (Finn et al. 2015; Xie et al. 2018; Salehi et al. 2020). Some past work has taken these results to suggest that measuring FC across different states can be a way to maximize individual identification (i.e., fingerprinting) across people (Finn et al. 2015). Here we take the complementary view that by studying single individuals, we may be able to provide a more accurate depiction of how brain networks are altered during task states. That is, a precision fMRI approach may reveal novel idiosyncratic components of how brain areas interact during complex tasks, which will necessarily be masked in approaches based on looking only for commonalities across people.

The goal of this paper is to determine the extent to which task state prediction is affected by individual differences. That is, how much accuracy exactly does one lose by trying to predict task states across people? What does this tell us about the individual selectivity of network features during tasks? This manuscript represents a deep dive into quantifying these prediction effects and their impact.

To address these questions, we used the Midnight Scan Club (MSC) precision fMRI dataset – which contains over 10 hrs of fMRI data per participant - to build classifiers to discriminate brain states (task vs rest) from a single person’s multi-session data. We then tested whether these classifiers could accurately determine task state in independent data from either the same person or a different person. Our results demonstrate that individualized classifiers can predict task state with high accuracy. While the classifiers do generalize to new subjects, they also exhibit sensitivity to person-specific features that are obscured in cross-subject data. This work shines a light on previously masked – individual-specific – characteristics of how functional brain networks are altered during task states.

## METHODS

### Overview

Our goal in this project was to investigate whether there are individual differences in how brain networks are altered during diverse task states. We used the precision fMRI Midnight Scan Club (MSC) dataset for this analysis (Gordon, Laumann, Gilmore, et al. 2017). The MSC dataset and its processing have been described extensively in (Gordon, Laumann, Gilmore, et al. 2017; Gratton et al. 2018), and we include a summary of the components relevant to this study below. For some replication analyses, we used data from an independent dataset recently collected at Northwestern University: the Individual Brain Networks Study (iNetworks) dataset. We describe relevant acquisition, task, and processing details of that dataset in the Supplemental Methods.

For both datasets, we created functional connectivity matrices from task and rest data for each participant, task, and session. We then used a machine learning approach to train classifiers to distinguish rest from task on a subset of sessions and then tested these classifiers on held out data from either the same participant or a new participant. Variations on this analysis were conducted with different classifiers, tasks, and feature subsets, as described below.

### Midnight Scan Club dataset

The MSC dataset includes data from 10 individuals (5 females, ages 24-34) with ten fMRI sessions each. Each session occurred on a separate day, beginning at midnight. Sessions were completed within 7 weeks for all participants. Each session included fMRI data for a resting-state scan and 4 tasks (see Task Designs and Analysis). Participants provided informed consent and procedures were approved by Washington University Institutional Review Board and School of Medicine Human Studies Committee. Participant MSC08 was excluded from the study due to high levels of head motion and sleep during rest (more details in (Gordon, Laumann, Gilmore, et al. 2017)). MSC09 was also excluded due to insufficient samples in the motor task (only 5 of the sessions had sufficient low motion data, see *Functional Connectivity Processing*). This resulted in eight participants used in the final analyses.

### MRI Acquisition

MRI data were acquired on a 3T Siemens Trio. Four T1-weighted images (sagittal, 224 slices, 0.8 mm isotropic resolution, TE = 3.74 ms, TR = 2.4s, TI = 1.0s, flip angle = 8 degrees) and four T2-weighted images (sagittal, 224 slices, 0.8 mm isotropic resolution, TE = 479 ms, TR = 3.2s) were collected for each participant across two separate days. Functional MRI data was collected using a gradient-echo EPI BOLD sequence (TE = 27ms, TR = 2.2 s, flip angle = 90, voxels = isotropic 4mm^3^, 36 axial slices) during each of 10 sessions. The same sequence was used for task and resting-state data. The same parameters were also used to collect a gradient echo field map acquired during each session for de-warping of the functional data.

### Task Designs and Analysis

Functional MRI data were collected during five conditions described briefly below (for a more thorough description see (Gordon, Laumann, Gilmore, et al. 2017; Gratton et al. 2018)). Task activations were modeled with a generalized linear model (GLM) using in-house software written in IDL (Research Systems, Inc.). GLM residuals were used for time-series correlations, following a background connectivity approach (Fair et al. 2007; Al-Aidroos et al. 2012). Task blocks/runs of the same condition were concatenated together for each session prior to creation of FC matrices.

#### Resting-state

Each session started with a 30 min resting state scan, wherein participants were asked to stare at a fixation on the center of a black screen.

#### Motor Task

The motor task was adapted from the Human Connectome Project (Barch et al. 2013) and consisted of a block design in which participants are told to move either left or right hand, left or right foot, or tongue based on visual cues. This included two runs in each session (7.8 min). Each block began with a 2.2s cue, followed by a fixation caret flashing every 1.1s to signal a movement. Each run included two blocks for each type of movement and three fixation blocks (15.4s). For this GLM each motor condition was modeled separately with a block regressor convolved with a hemodynamic response function.

#### Semantic Task

Each session included two runs (14.2 min) of a mixed block/event-related design modeled on tasks in (Dubis et al. 2016). This included four blocks per run, two for semantic, two for coherence (see below for coherence). The semantic task was a verbal discrimination task where participants were visually presented with short words and asked to identify if they were nouns or verbs (50% nouns, 50% verbs). Task blocks began with a 2.2s cue indicating which task was to be conducted in the following block; after this 30 individual trials were presented. Trials consisted of words presented for 0.5s with jittered 1.7-8.3s intervals. Participants responded to whether each word was a verb or a noun. After, participants were presented with a cue (2.2s) indicating the end of a block, with 44s fixation periods separating each block. The semantic and coherence tasks were modeled together in a single mixed block/event-related GLM. Separate regressors were included for each task block and for events (start and end cues in each task, correct and incorrect trials of different types). Events were modeled with delta functions for 8 separate time points to model the time course of responses using an FIR approach (Ollinger et al. 2001).

#### Coherence Task

Coherence task blocks were interleaved with the semantic task in the same mixed runs and followed the same timing structure and analysis as the semantic task. Individual trials consisted of arrays of Glass-like patterns, with white dots on a black screen that varied in arrangement (0% or 50% coherence to a concentric arrangement, both presented with 50% frequency) (Glass 1969). Participants were told to identify dot patterns as concentric or random.

#### Memory Task

The incidental memory task employed an event-related design with 3 runs per session (15 min). A single run was collected for each type of stimulus (faces, scenes, words). During trials of the run, participants made binary decisions to categorize stimuli (male/female for faces, indoor/outdoor for scenes, abstract/concrete for words). In each run, participants viewed 24 images repeated 3 times, making this task a test of implicit memory (note that the repeats were orthogonal to the participants’ task goals). Images were presented for 1.7s, with jittered 0.5 – 4.9s intervals. For the memory GLM, trials were modeled based on stimulus type and number of repetitions with delta functions across 8 time points.

### Structural MRI Processing and Surface Registration

The procedure used to process the structural MRI data and align the data with the surface can be found in extensive detail elsewhere (Laumann et al. 2015; Gordon, Laumann, Gilmore, et al. 2017). The high-resolution structural T1 images were first aligned and averaged together, then registered to a volumetric Talairach atlas using an affine transform. These averaged template T1s were used to generate a cortical surface in Freesurfer (Dale et al. 1999), which was hand-edited to improve surface accuracy. The native freesurfer surface was registered to fs_LR_32k space using a similar procedure described in (Glasser et al. 2013).

### Functional MRI Pre-processing

All fMRI data first underwent pre-processing in the volume to correct for artifacts and align data. This included slice-timing correction, frame-to-frame alignment for motion correction, and intensity normalization to mode 1000. Functional data was registered to the T2 image, which was registered to the high-resolution T1 that had been registered to template space. Functional data underwent distortion correction (see (Gordon, Laumann, Gilmore, et al. 2017) for more details). Registration, atlas transformation, resampling to 3mm isotropic resolution, and distortion correction were combined and applied in a single transformation (Smith et al. 2004). All remaining operations were completed on the atlas transformed and resampled data.

Task fMRI data were processed in the volume using a GLM, with the approach and regressors described above (*Task Designs and Analysis*). The residuals from this model were used to compute task functional connectivity (as in (Gratton et al. 2018)), following the same steps as rest (described below).

### Functional Connectivity Processing

To reduce artifacts, data underwent initial functional connectivity (FC) processing in the volume, with methods described in detail in (Power et al. 2014; Gordon, Laumann, Gilmore, et al. 2017). This involved demeaning and detrending the data, nuisance regression of motion parameters and signals from white matter, cerebrospinal fluid, and global signal, as well as their derivatives. High motion frames (framewise displacement, (Power et al. 2012) FD > .2mm along with sequences containing less than 5 contiguous low motion frames, the first 30s of each run, and runs with <50 low motion frames) were censored and replaced with power spectral matched data. For participants MSC03 and MSC10, motion parameters were low-pass filtered (<0.1Hz) before FD calculation. Following motion censoring, bandpass filtering (.009Hz-.08Hz) was applied. Cortical functional data was then registered to the surface and smoothed (Gaussian kernel, sigma=2.55mm) with 2D geodesic smoothing on the surface and 3D Euclidean smoothing for subcortical volumetric data (see (Gordon, Laumann, Gilmore, et al. 2017) for more details). Finally, interpolated high motion frames were removed before functional connectivity analysis. To be included, each run was required to include at least 25 low-motion volumes, and each session required at least 50 volumes total per task; we removed 5 sessions based on these criteria. Task data was limited to task periods for each run (excluding fixation frames).

### Functional Networks

#### Regions and Systems

This study examined functional connectivity among 333 cortical parcels defined based on boundary-mapping techniques in a large group of independent participants (Gordon et al. 2016). These 333 parcels are divided into 13 functional systems: somatomotor (SM), somatomotor lateral (SM-lat), visual (Vis), auditory (Aud), cingulo-opercular (CO), salience (Sal), frontoparietal (FP), dorsal attention (DAN), ventral attention (VAN), default mode (DMN), parietal memory (PMN), and retrosplenial (RSP). Low signal regions that grouped poorly into a system were put in an ‘‘unassigned’’ group.

#### Functional Connectivity (FC)

FC was computed by averaging the BOLD time course within each parcel, after removing censored and interpolated frames, and computing linear correlations between the time-series of each pair of parcels. Task data were limited to task periods within each run (i.e., excluding fixation periods). FC values were Fisher transformed for normality. FC was represented with a parcel by parcel functional network matrix, sorted by system edges (FC values for a particular region-to-region pair) along the diagonal blocks represent within-system correlations, and edges in the off-diagonal blocks represent between-system correlations. Edges taken from the upper triangle of this matrix represent all unique pairwise parcel relationships resulting in 55,278 values for each task of a given session. Unique edges were used as features in the machine learning classifier.

### Machine Learning

We employed a machine learning approach at the individual level where models were trained to classify task state using data from a single participant, which we call individualized classifiers. Models were trained and tested on independent data using a leave-one-session-out cross-validation framework, iteratively leaving out data (both task and rest) from an entire session for training and then testing on either this left out data or data from a single session in another person (for schematic see **Fig. 1**). Classification was performed using different classifiers, subsets of tasks, and subsets of features as specified below.

**Figure 1.**
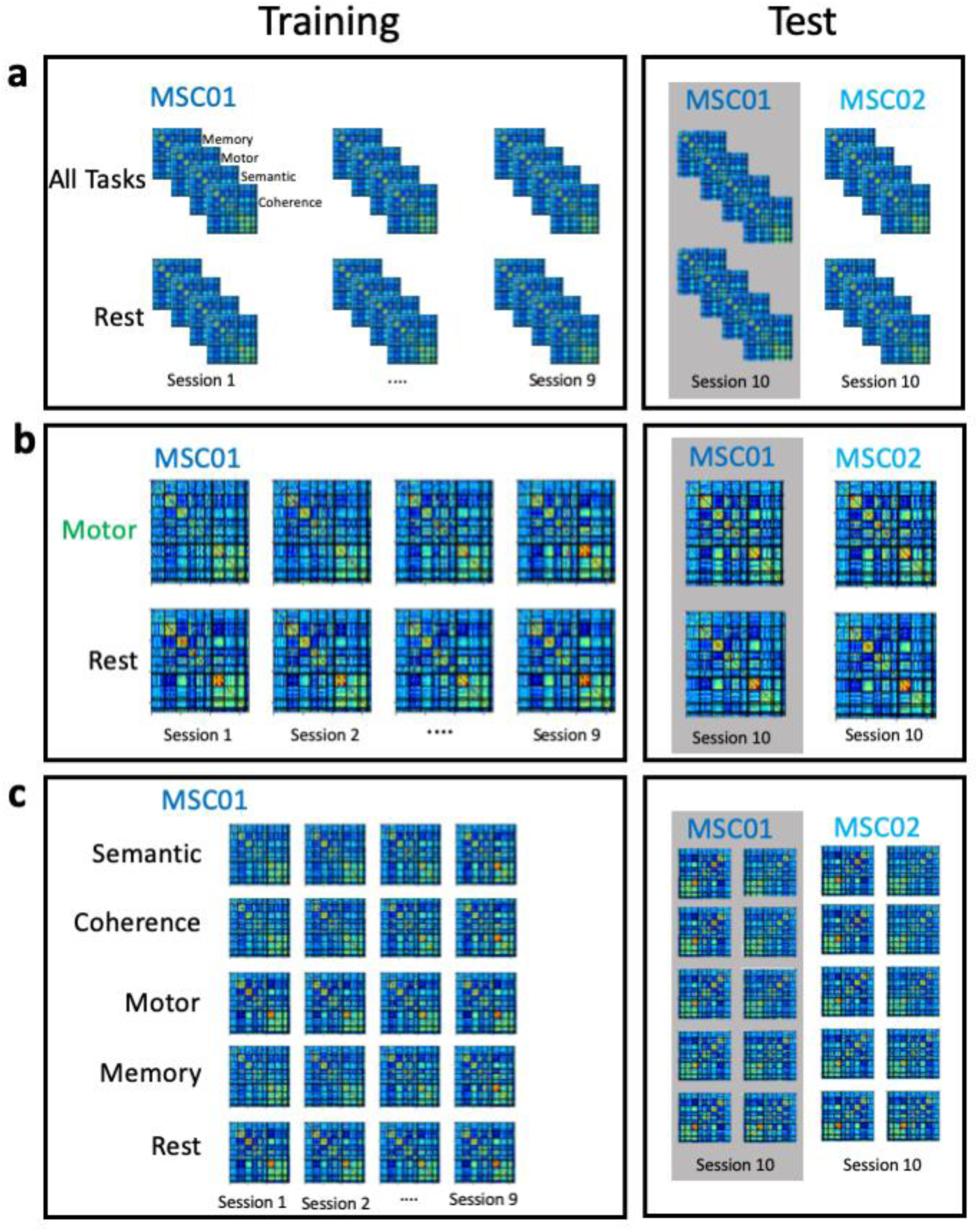
Overview of each analysis. Classifiers were trained to distinguish task from rest functional connectivity using many sessions of data from a single participant with ridge regression. **(a**) For our first set of models, classifiers were trained to discriminate between all tasks and rest (80 total samples per participant). **(b**) In the second group of models, classifiers were trained to distinguish a single task (vs. rest) (**c**) Finally, multiclass models were trained to discriminate among all task states at once In all cases, models were then tested on independent session data from either the same person (gray bar) or a different person; performance was contrasted across these tests to identify generalizable or individually-specific components of task state.

#### Ridge Regression Classification

We used ridge regression for classification due to the high collinearity among FC edges and the relatively small number of samples. Ridge regression implements a shrinkage estimator that can improve performance in the presence of multicollinearity (Fitrianto and Yik 2014). This pipeline was implemented using Python’s Scikit-learn package (Pedregosa et al. 2011). As a control, we also conducted analyses using support vector classification and logistic regression and found that all models resulted in similar patterns of performance across conditions (see **Fig. S1**).

#### Binary classification of task state across people

Our first goal was to determine the extent to which a model trained on one participant would be able to differentiate task from rest both in that person and in new people.

#### Training and testing procedure

We conducted two versions of binary classification to examine task state prediction from FC data. The “<u>*All Task*</u>” classifier was trained to distinguish all tasks (memory, motor, semantic, coherence) from rest in a single binary classifier. This classifier was used as a higher sample test of the ability for classifiers to distinguish task states from rest. This classifier had 80 total samples per participant: 40 task samples (10 sessions of motor, coherence, semantic, and memory tasks) paired with 40 rest samples (rest split into four 7 min increments for each of 10 sessions; we split rest samples in order to balance our training set and avoid biasing the model towards the task samples). For each individual, we then trained a classifier to predict task state (task vs rest) based on FC using a leave-one-session-out cross-validation scheme as described above (90% training data, 10% testing per fold). Each model was also tested on a single session from other participants (resulting in 7 “other person” tests per trained model). This training/testing process was repeated for all folds and all participants.

The second “<u>*Single Task*</u>” binary classifier classified a single task at a time (e.g., memory vs. rest). This classifier allowed us to determine whether the All Task results were consistent for each specific task condition. The same procedure was used for the “All Task” classifier, except in this case each classifier was based on a total of 20 samples per participant (10 from a single task, 10 from rest). Models were again trained on 90% of the data (9 sessions) and tested on either left out data from that same participant (1 session) or another participant. The procedure of training/testing an individual classifier was repeated for all 4 tasks (motor, memory, coherence, semantic).

#### Model evaluation

Classifier accuracy (total percentage of test samples classified correctly), task predictive value (TPV; correct labeling of task divided by any labeling of task), and rest predictive value (RPV; correct labeling of rest divided by any labeling of rest) were used to evaluate model performance for each individualized classifier. For each classifier, test performance was averaged over folds. Data is presented separately for each participant (each individualized classifier), representing an 8-fold replication of model training and testing on independent people. Performance values were contrasted with permuted null models (see below) to determine significance.

#### Model significance

First, we tested whether the performance of each classifier was significantly greater than chance when compared to a permuted null model. For each model, we trained a null classifier using the same FC data, but permuted how samples were labeled as task or rest. The null classifiers were subsequently tested on held-out data using the same cross-validation framework described above for each type of classifier. We repeated this process 1,000 times for each task, averaging permuted and true accuracies across all individualized classifiers to compute an omnibus statistic. We then calculated permutation-based p-values by comparing the average true classification accuracy relative to the distribution of classifier accuracy of the null classifiers (number of permutations that exceeded true model performance+1/1000+1). Additional tests were run per individualized classifier, comparing that classifier to a permuted null sample.

Second, we tested whether the performance of a classifier significantly differed when applied to independent data from the same individual vs. a different individual. We again used a permutation approach, in this case randomly shuffling the same-person or different-person labels of the test sets, then calculating the difference in accuracy scores. We repeated this process 1,000 times to create a null distribution, after which the true difference was contrasted with the null distribution and used to estimate a permutation-based p-value.

#### Groupwise cross-subject models

To evaluate how these measures compare to a standard machine learning approach we conducted a leave-one-*subject*-out analysis in which we trained the classifier using a single session of data across all participants in the dataset and tested on the left-out participant (training 7 participants, testing 1 participant on the same session; termed the ‘groupwise’ classifier). We then repeated this process across all participants and across all sessions. We conducted cross-subject models both for decoding all tasks from rest (as in the *All Task Analysis*) and models for discriminating a single task from rest (as the *Single Task Analysis*). Model performance is reported as the average accuracy across all folds. Note that to fairly compare this groupwise model to the individualized models, we created separate individualized models using the same number of samples as the groupwise approach (e.g., 7 sessions for training, 1 session for testing in the single task, etc.); these were used for comparison to the groupwise classifier.

We also conducted a second form of groupwise analysis in which all possible within-person data was used for model training (*Groupwise Max* e.g., training on 7 participants using all 10 sessions per participant, testing on 1 participant with 10 sessions of data); note this model includes substantially more training data than the individualized models. Both of these groupwise classifier models (*Groupwise* and *Groupwise Max*) were used in our analysis examining the similarity of feature weights across the different approaches (see below). Finally, we conducted a groupwise analysis in which we iteratively included additional sessions in the training set from 1 session to 10 sessions per participant (7-70 task:rest sample pairs) in order to determine how groupwise classifier performance was affected by increasing amounts of data.

#### Binary classification of similar task conditions

To determine if these effects would remain significant when predicting across highly similar task conditions we trained a binary classifier to discriminate between different conditions of the memory task runs with the first, second, or third repetition of presented stimuli (each participant performed the same categorization judgment at each repetition). This classification procedure followed the same procedure as described earlier in which we trained a classifier using leave-one-session-out cross-validation for a given participant, then tested on the same or a different participant including model evaluation and significance.

#### Multiclass classification of task state across people

We next asked if finer-scale discriminations of task state could also be decoded from FC. To address this question we trained a *Multiclass* classifier to discriminate among all 5 of the separate states at once (rest, motor, memory, semantic, coherence; chance = 20%). We then repeated the same procedure as above for cross-validation (leave-one-session-out for a given participant, then test on the same or a different participant) and model evaluation and significance. For this analysis, we also created confusion matrices of the test performance to better assess errors.

#### Classification of models built on individualized network maps

We conducted an additional set of classification analyses to evaluate whether differences in participant performance could be attributed to differences in underlying spatial layout of large-scale networks. To address this question, we reconducted our classification based on individually defined network layouts. Previously published (Gordon, Laumann, Gilmore, et al. 2017) individually defined parcels and network assignments for the MSC dataset were used for this analysis. These parcels and networks were defined independently for each participant using data-driven procedures. Importantly, previous results show that these individual network assignments reach high reliability with the precision fMRI data employed here (Gordon, Laumann, Gilmore, et al. 2017). We then calculated functional connectivity among each of these individually defined regions for each state. Finally, we averaged across each of 14 networks that could be identified consistently in each individual (default mode network, visual, frontoparietal, dorsal attention network, premotor network, ventral attention network, salience, cingulo-opercular network, somatomotor dorsal, somatomotor ventral, auditory, parietal memory network, and retrosplenial network; as in (Gratton et al. 2018) Supp. Fig. S7). This resulted in FC matrices for each person, task, and session with a 14 × 14 network dimensionality – notably while the dimensionality and network labeling was matched across participants, each person’s networks were defined separately to optimize accounting for that person’s specific spatial layout of networks. We then extracted the features from these matrices and conducted the same set of individualized model training and testing as described in the binary classification section above. For comparison, we also conducted a similar analysis, averaging across each of 13 networks that could be identified at the group level. This resulted in FC matrices for each person, task, and session while matching across each person’s network definitions.

### Feature Analyses

Next, we asked if the classification of task state was dependent on specific features or networks. We took two distinct approaches to address this question.

#### Feature weight analysis

First, we examined the feature weights from our individualized classifiers. To do this we extracted the feature weights for the *All Task* binary classifier (see above) averaged for each region (row of the correlation matrix; averages were taken on the absolute values of the original weights). We qualitatively examined the variability of average feature weights across folds to look at consistency within a person. We then examined the variability of average feature weights across individuals (in this case averaged over folds) to look at the consistency of feature weights across people. We also conducted the same procedure when analyzing feature weights at the single task level. To evaluate whether feature weights are degraded or fundamentally different when using standard groupwise approaches we conducted the same procedure on the Groupwise approach for the *All Task* binary classifier and the single task classifiers. To compare features across different models, we took Pearson correlations across each classifier’s feature weights at the fold level.

#### Feature selection analysis

Secondly, we conducted additional quantitative classification analyses with only subsets of features to determine which features were sufficient for classification. Three forms of feature selection were performed: (1) features were selected from specific blocks of network to network connections (e.g., all of the features associated with connections between the frontoparietal network and the default mode network). Due to the low feature size associated with specific blocks, we standardized the data by removing the mean and scaling to unit variance of the training set. We then transformed all test sets using the mean and standard deviation from the training set (Pedregosa et al. 2011). (2) To examine properties for full networks, features were also selected for entire network rows (e.g., FC in all matrix rows associated with default mode network regions), and finally (3) as a comparison, random features numbering 10 – 50,000 were randomly selected from the full matrix (this process was repeated 1,000 times). Model training and testing was then conducted as described in the binary classification section for both the *All Task* and *Single Task* analyses. This process was repeated independently for each feature subset and for each separate participant.

### Data Quantity Dependence

Since data quantity has been demonstrated in the past to affect the reliability of FC matrices (Laumann et al. 2015; Noble et al. 2017), we examined whether model performance depended on sample number (i.e., the number of FC matrix pairs which were included in the training data). To determine how the number of samples in the training set influenced model performance, we re-built the *All Tasks* classifier described above using varying amounts of sessions from a single individual. Beginning with 80 samples of FC matrices (40 task and 40 rest) for each person, we then sub-selected between 16 – 80 samples (in matched task/rest pairs) and repeated model training and classification, using a leave-one-session-out cross-validation approach. We also tested the model on each of the other participants’ task and rest data. This training/testing process was repeated for all folds and all participants. This process was repeated 1,000 times for different random pairs of samples per individualized classifier.

### Replication from Independent Dataset

To determine the extent of our findings, we used a similar precision fMRI dataset (as described in Supplementary Methods). In which we trained an individualized classifier using a single person’s multi-session data. Due to the number of available sessions we trained a classifier using a leave-one-session-out approach maintaining each run as a separate FC matrices (e.g., training on 3 sessions with 12 runs of mixed and 12 runs of rest to match, testing on 1 session with 4 runs of mixed and 4 runs of rest). We conducted the same approaches as described above in which we trained classifiers using all task data vs rest (binary), all task data and rest (multiclass), and single task data vs rest. In addition we conducted a similar groupwise approach for each condition in which we randomly sampled the number of subjects to match the number of samples trained in each individualized approach.

## Data and Code Availability

Midnight Scan Club data is publicly available (https://openneuro.org/datasets/ds000224). Code for analysis related to MSC pre-processing can be found at https://github.com/MidnightScanClub. Code related to the analysis in this paper will be located at https://github.com/GrattonLab/Porteretal_taskprediction.

## RESULTS

In this project, our goal was to improve our understanding of how brain networks are altered during tasks by using an individual-level approach to test whether (and which) state changes in functional networks generalize or are specific to particular individuals. We measured functional brain networks across 4 task states and rest in highly sampled individuals with 10 separate fMRI sessions from the MSC dataset (Gordon, Laumann, Gilmore, et al. 2017). We used a machine learning approach, wherein we trained classifiers to predict task state based on data from a single individual. We then tested whether these classifiers could effectively classify state on independent data from the same individual or data from other individuals (**Fig. 1**): accuracy differences between these test cases were used to identify generalizable and individually-specific components of task state. Eight individuals were examined, serving as 8 replications of our findings. Results were further confirmed through replication in 22 individuals in a novel independent dataset.

### Individualized classifiers can predict task state both within and between individuals

We first evaluated the performance of models trained in a single person to distinguish between task states and rest (binary: any task vs. rest) using independent data in the same individual. We began by training a classifier using ridge-regression with leave-one-session-out cross-validation (80 samples per person, 72 training, and 8 testing per fold; see *Methods*). In all participants, models performed well at discriminating task from rest for new data from that same person (within-person test: M=.97+/−.01; p<.001 in contrast to a permuted label null). Models also predicted task state significantly greater than chance when tested in other participants, indicating they were able to generalize (between-person test: M=.68+/−.11; p<.001 for all participants; **Fig. 2a;** see **Fig. S2** for results and p-values per person).

**Figure 2.**
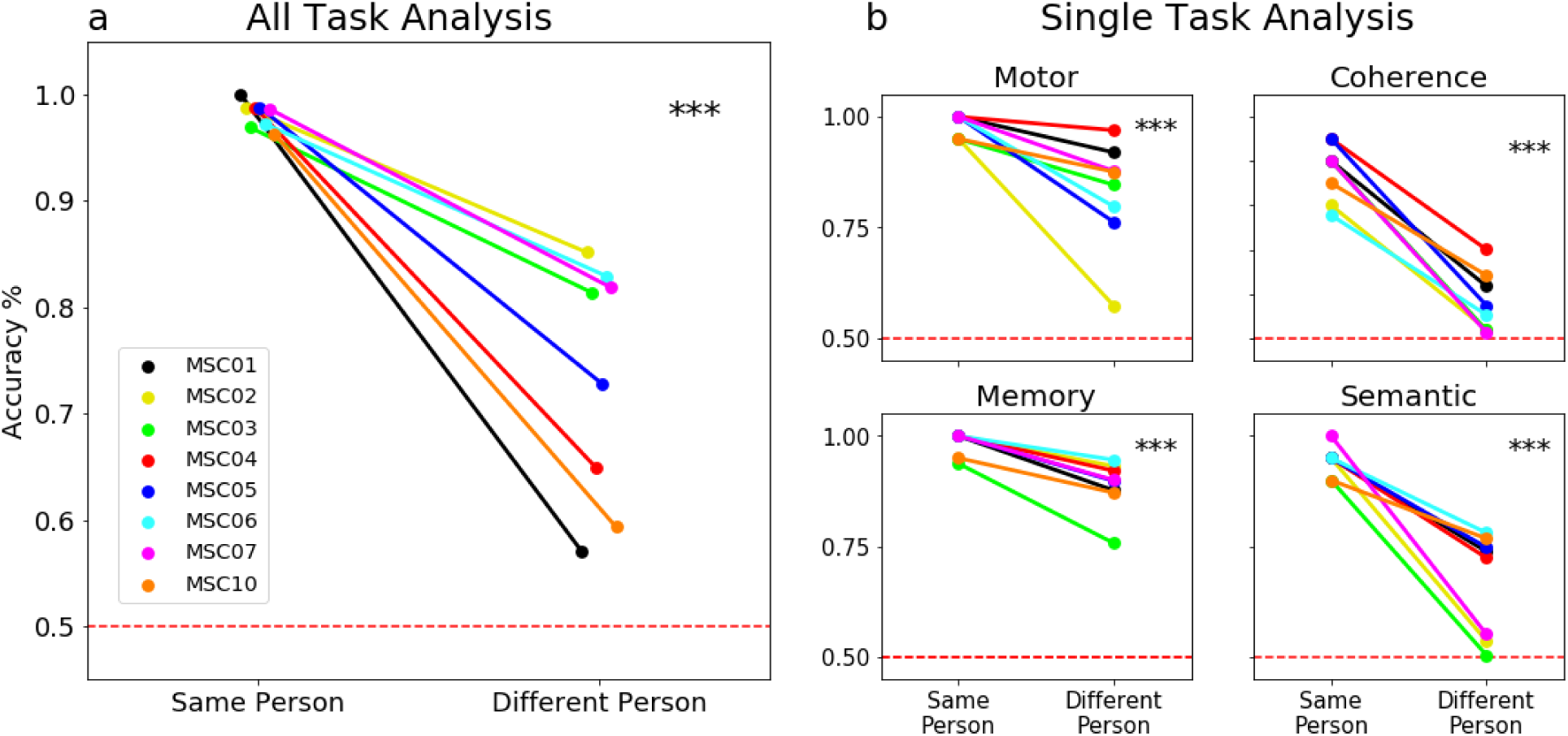
Differences in classifier accuracy within and between-person. **(a)** Performance of an individualized machine learning classifier for discriminating task from rest states when tested on independent data from either the same individual or different individuals (colored lines = different participants). Initial models contrasted all task states with equal samples of rest (N = 80 samples total per person). **(b)** Secondary analyses trained models to discriminate a single task from rest. On average, all models were able to predict task state significantly better than chance in the same person (p<0.001 when compared to a permuted null with randomized task/rest labels) and in new people (p<0.001; red dashed lines = chance). However, model performance was significantly higher when tested on data from the same participant relative to other participants (*** p<0.001 compared to a permuted null on within/between person labels; All Tasks vs. Rest: effect size=.24; Motor vs. Rest=.15; Coherence vs. Rest=.30; Memory vs. Rest=.09; Semantic vs. Rest=.27;). Similar results were seen when tested with other machine learning algorithms (***Fig. S1***). For a more detailed breakdown of person-to-person performance see ***Fig. S2)***.

We then evaluated prediction performance for specific tasks. Again, classifiers were trained on a subset of a given participant’s data and tested on data from an independent session from the same person or another person using leave-one-session-out cross-validation. We saw consistent results across tasks and participants (**Fig. 2b**), with significant task state decoding when tested on data from both the same (within-person test: Coherence M = .87+/−.06; Memory M = .99+/−.01; Motor M = .98+/−.02; Semantic M = .94+/−.03; all p<.001) and other individuals (between-person test: Memory = .89+/−.1; Motor M = .78+/−.15; Semantic: M = .66+/−.14; and Coherence: M = .57+/−.1; all p < .001). These results demonstrate that functional networks carry task state information, which can be used to make accurate predictions about state in the same and other participants.

### Task state prediction is higher within than between individuals

Although we were able to predict task state both within and between individuals, task prediction performance differed significantly in these two test sets. Classification was significantly more accurate when discriminating between task and rest within the same person vs. other individuals (mean effect size=.24, p<0.001 based on permutation, see *Methods*; **Fig. 2a**). These findings were robust to the number of samples included in the training set (**Fig. S3**, when testing on 16-80 samples): classification of task state generally improved for both the same and other people with more samples, but more dramatic improvements were seen when tested in the same person. This effect remained consistent when predicting tasks separately (**Fig. 2b**). Within-person classification was significantly higher than between-person classification for all comparisons (mean difference of within and between-person performance: motor=.15; coherence=.30; memory=.09; semantic=.27, p<.001 for each separate comparison). In all cases, within-person classification was higher than between-person classification, indicating that while classifiers could generalize, task states exhibit a substantial amount of individual-specific characteristics.

Notably, the between-person test performance was similar to the performance seen for a more classic leave-one-*subject*-out (groupwise) classifier (where training was performed on a session from many different participants and tested on a novel participant; **Fig. S4**). In all cases, groupwise classification performed similarly to the between-person test sets and worse than the within-subject classifiers (mean accuracy difference of within-person compared to groupwise classification for all tasks =.17; single task analysis: motor = .15, coherence = .29, memory = .09, semantic = .25). We found that increasing the number of training sets in the groupwise analysis improved model performance, eventually reaching levels in the range of that seen with individualized classifiers. However, this generally required over 70 samples of training data (achieved, given the nature of our dataset, by adding additional sessions of data from each participant), nearly 4 times as much training data compared to the individualized classifiers performing at similar levels. These results suggest that groupwise classifiers can achieve good accuracy, but require more training data to do so – and interestingly suggest that addition of more single-subject data can also lead to a more robust measure of shared features across people.

Moreover, we find that this effect is consistent even when training a classifier to decode very similar task conditions (**Fig. S5**). The memory task contained three separate runs in which the same stimuli were repeated; during each run, participants saw the same stimuli and completed the same task (a categorization judgment orthogonal to the repetitions). Thus, in each case, the sensory, motor, and explicit task demands were matched, with the only differences arising from the number of previous exposures to the stimulus. We then asked whether individualized functional connectivity classifiers could discriminate very similar conditions such as these. We trained a classifier along a single person’s multi-session data to discriminate among the three conditions in all possible pairs (e.g., presentation 1 vs. presentation 2), and then tested this ‘similar condition’ classifier on left out data from either the same person or a new person. All classifiers were able to predict performance better than chance (**Fig. S5**, p<.001). Consistent with the previous results, however, we found that classifiers achieved significantly higher accuracy when tested in the same person than new people (Mean difference of within and between person performance accuracy = .15, p<.001).

The significantly higher performance of classifiers tested on the same individual across each of these tests suggests that there are substantial task network effects that are unique to individuals and do not generalize between people. These findings reveal that individualized, within-person analyses unmask substantial information on how brain areas communicate during tasks.

### Within-subject classifiers exhibit fewer prediction biases than between-subject classifiers

To provide a more thorough investigation of classification accuracy, we calculated task predictive value (TPV; correct labeling of task divided by any labeling of task) and rest predictive value (RPV; correct labeling of rest divided by any labeling of rest). When tested within the same individual, TPV and RPV scores were both consistently high (all tasks vs. rest, within-person: TPV=.97+/−.01, RPV= .99+/−.006; single tasks vs. rest: see **Table S1**). More errors were seen for between-person analyses (all tasks vs. rest, between person: TPV= .69+/−.1, RPV= .93+/−.07; single tasks vs. rest: see **Table S1**). The errors in state classification for new participants differed by classification procedure: in the all task vs. rest discrimination, rest was frequently misclassified as a task, whereas in the single-task analysis certain tasks (coherence, semantic) were frequently misclassified as rest. Importantly, in all cases, the within-person analyses had both high TPV and RPV. These results suggest that classification across individuals can result in biased errors dependent on the training set. Within-person classification is less subject to these biases.

### Individualized task state prediction is high across many tasks at once

Next we asked if individualized task state prediction could extend beyond a binary classification (task vs. rest) to classify varied task states at once. Using ridge regression we trained a multiclass model on data from a single individual to distinguish between the 5 different task states (rest, motor, memory, semantic, coherence) and then tested this model on states from an independent session either from the same person or a new person (**Fig. 3a**). In all participants, multiclass models were able to predict task state above chance for both within and between-person test sets (Within person M=.83+/−.05, p<.001; Between person M=.53+/−.07, p<.001 relative to null permutation model). Again, however, we found that within-person classification significantly outperformed classification across individuals (mean effect size= 0.3, p<.001 based on permutation). An analysis of the errors in multiclass performance showed greater overall errors in between-person classification and a greater propensity to classify other tasks as rest (especially for the motor and semantic tasks, see **Fig. 3**). Thus, individualized classifiers can perform even multiclass predictions with good accuracy, substantially out-performing more standard between-person approaches. These results suggest that individualized classifiers are able to capitalize on idiosyncratic task state information and may be useful in providing robust analyses of the commonalities and differences across task states.

**Figure 3.**
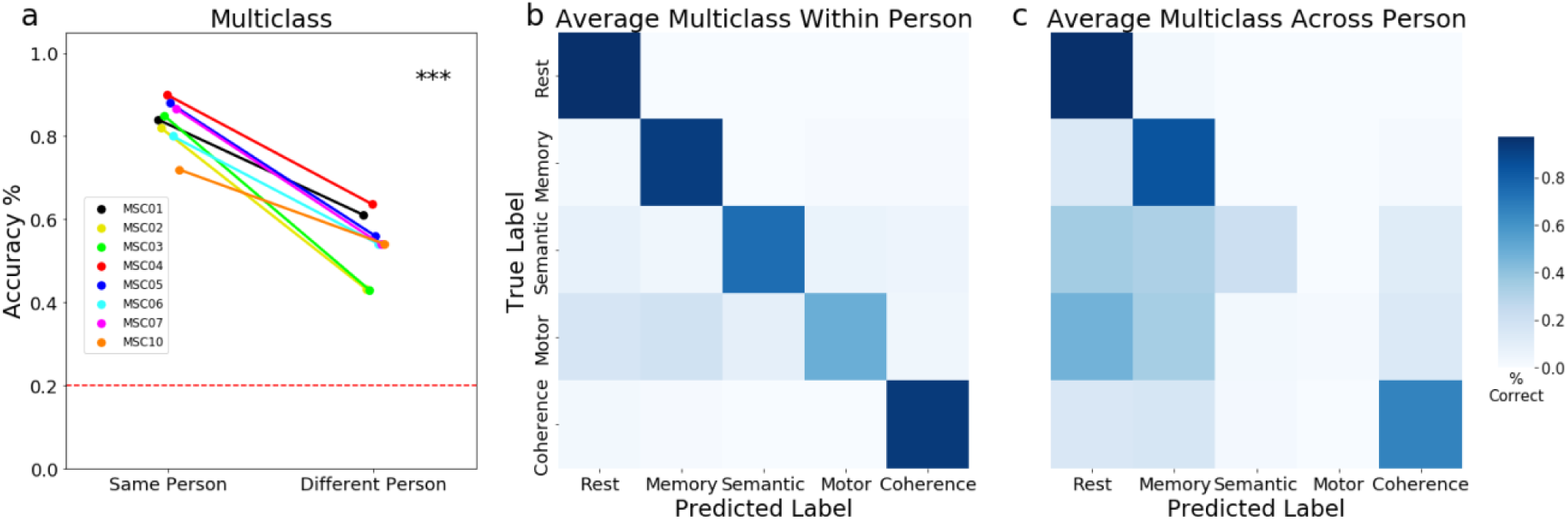
Multiclass Performance. (a) Here we trained models to discriminate all tasks and rest using a multiclass approach. For a detailed breakdown of performance in each individualized multiclass model see ***Fig. S6***. Multiclass prediction was significant both in the same and new people (*red line = chance performance across 5 states)*, with higher performance in the same individual (*\*\*\* p<0*.*001)*. In addition we calculated confusion matrices for classifiers tested on independent data from the (**b**) same or (**c**) a different person. Classifiers performed relatively accurately when classifying new data in the same person, exhibiting few biased errors. In contrast, classifiers tested in new people exhibited more errors, especially for the motor and semantic tasks. Classifier performance is averaged across participants (see Fig. S6 for confusion matrices from individual participants).

### Task state features are robust but distinct across individualized and group classifiers

Next, we asked which brain network connections are most important for predicting task states. We addressed this question through complementary analyses on feature weights and feature selection procedures. First, we examined the anatomical distribution of the average feature weights for each brain region to better understand which features contributed the most to task state decoding. Feature weights were fairly consistent across folds for a given individualized classifier (**Fig. 4a**). However, strong feature weights mapped onto different regions across people (**Fig. 4c**), likely underlying the variation in classification performance across people. Feature weights also varied somewhat across tasks in the separate single task classifiers (**Fig. S7**), indicating the presence of task-specific components to prediction and likely underlying the success of multi-class prediction. A similar pattern of distributed and individually divergent feature weights was seen even for classification of very similar conditions from the same task (different repeats of stimuli from the memory task; **Fig. S13**), although feature weights appeared overall lower and less generally widespread than in the more general task comparisons, as might be expected given the added specificity of this contrast. Feature weights from more classic leave-one-subject-out groupwise classifiers appeared lower in magnitude and showed a distinct distribution relative to individualized classifiers in all cases.

**Figure 4.**
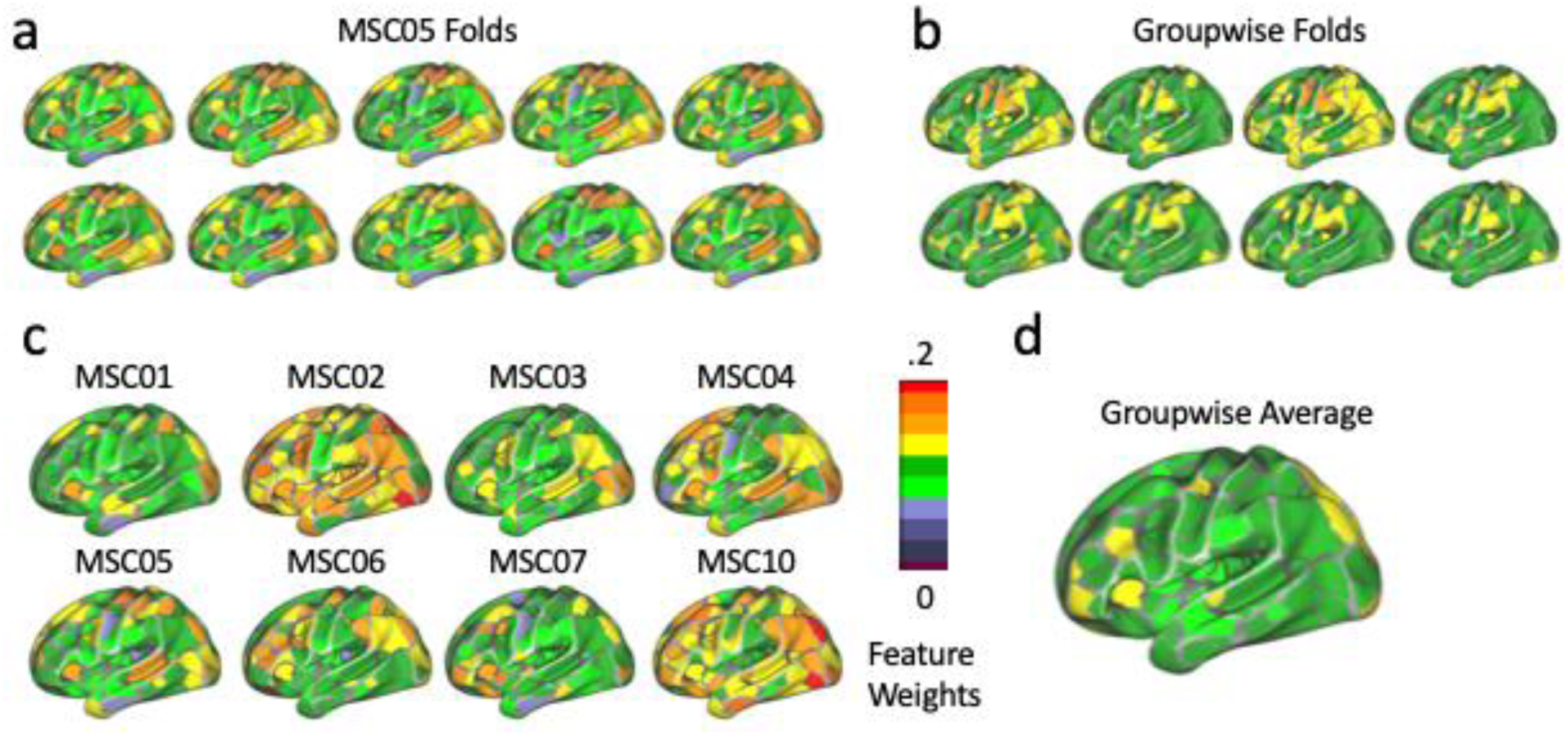
Feature Weight Analysis. Average absolute feature weights for each brain region for classifiers built to discriminate between all task states and rest. **(a)** Feature weights for a single person (MCS05) across each fold of the classifier. **(b)** Feature weights from the standard groupwise approach for each fold of the classifier (in this case, training data comes from different participants, rather than different sessions of the same participant). **(c)** Average feature weights for each individualized classifier for each participant. **(d)** Average feature weights for the groupwise approach. Note that while feature weights are fairly similar across folds within a person, they show variation across people and compared to the groupwise approach.

We quantified these differences by correlating feature weights across folds of the individualized and groupwise classifiers (Fig. S8). Consistent with the observations above, feature weights were quite similar across folds within a person (r = 0.92), but distinct across individuals (r = 0.25). Intriguingly, feature weights were also distinct between individualized classifiers and the more classic groupwise classifier (r = 0.16). When the maximally performing groupwise classifier was used (including 280 sample pairs, 7x as much as any of the other individual classifiers), we find a slightly higher correspondence to the individualized classifiers (r = 0.38), but this is still substantially lower than what is seen for a given participant across folds. However, separate folds of each groupwise classifier are relatively consistent with one another (r=0.79), although slightly less consistent than what is seen with the individualized classifiers. Jointly, these results suggest that individualized and group classifiers capture robust, but distinct features of task state.

### Task state can be decoded from single networks

Next we asked whether single networks would be sufficient for predicting task state in individualized classifiers. To address this question, we built classifiers restricted to features in specific blocks of network-to-network connections (**Fig. 5**). We found good performance for most classifiers based on single network blocks. For within-person analyses, average block performance was 83% (range 61%-95%). 99% of blocks were able to decode task state on their own above chance (SD=.008). In between-person analyses, average block performance was 57% (range 51%-68%). 95% of blocks were able to decode task state above chance (SD=.04). (SD=.04).

**Figure 5.**
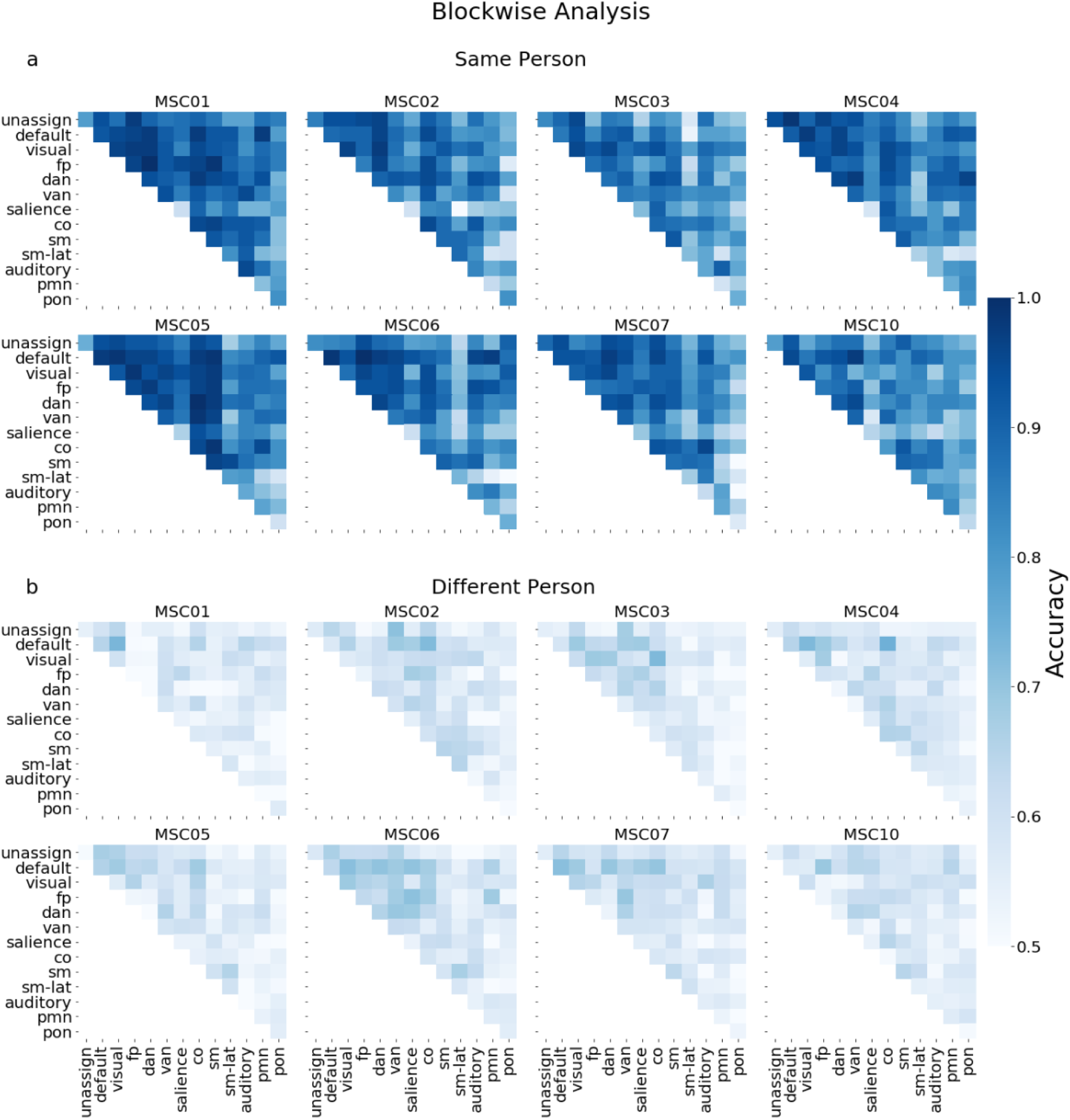
Single network block performance in decoding task state. Here we trained classifiers to decode all tasks from rest using subsets of network to network connections (each block was an independent classifier). We then tested the classifier on new data from either (a) the same person or (b) a new person. Due to low feature numbers prior to training the classifiers, we standardized features (see Methods). Task state information was distributed across many networks of the brain: in within-person analyses, 99% of blocks could decode task state; 95% of blocks could decode task state in between-person analyses. Within-person performance was consistently higher than between-person classification.

This sufficiency test indicates that task state classification features can be found in almost all networks. While performance levels varied by network block, these network blocks also differed substantially in size (different numbers of regions in each network lead to different numbers of features), and better performance was generally found for larger networks. To systematically test whether specific networks or simply higher feature numbers were important for classification performance, we contrasted the performance of features extracted from single networks (all connections associated with regions from a single network) relative to feature sets that were randomly selected from the full connectome. Random feature sets numbering 10-50,000 features were tested. In all cases, classifier performance improved with more features.

Moreover, at all feature numbers, within-person classifier performance outperformed between-person classifier performance (see **Fig. 6**). Interestingly, there were no cases where single networks performed significantly better than random feature selection when feature numbers were matched. Instead, in the majority of cases, single networks performed similar or worse than randomly selected features. This suggests that task state information, including individual-specific aspects of that information, is distributed across multiple networks that a classifier can utilize in prediction. We hypothesize that prediction is better with random features because random feature selection can utilize more independent sources of information from different networks (within network features are more likely to share correlated information, as has previously been found for age prediction performance (Nielsen et al 2020)).

**Figure 6.**
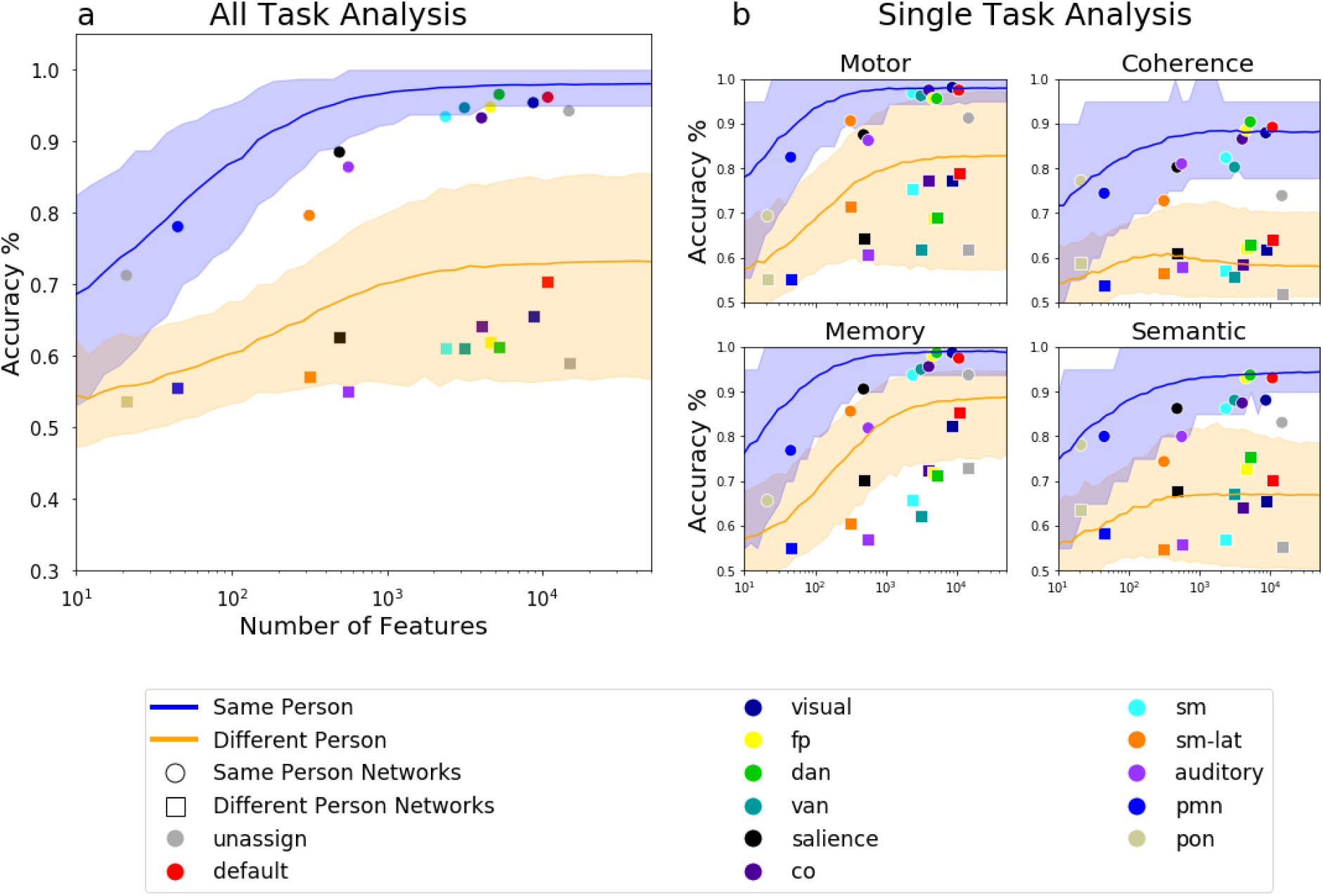
Overview of feature selection. Model classification performance when classifiers were trained on randomly selected features of different numbers, or features associated with a given network. Random feature number varied from 10-50,000, randomly indexed across all unique FC edges. Networks have been plotted on top at the position equivalent to their feature size. Accuracy is shown both for models tested on the same person (circle markers, blue line) or a different person (square markers, orange line). Error bars on the random feature selection line represent the 5th and 95th percentile across iterations. (**a**) Performance when training and testing to discriminate all tasks from rest. (**b)** Performance when training and testing to discriminate a single task from rest. For a breakdown of participants see **Fig. S9**.

### A within-person advantage is seen even with individually-mapped network regions

Most past machine learning studies rely on between-person analyses using a common standardized set of brain regions defined for group comparisons, which we demonstrate systematically underperform within-person models. This observation naturally raises questions about the source(s) of improved classification within-person. One possibility is that within-person classification is driven by differences in the underlying spatial positions of networks across individuals; within a person, these are relatively consistent over time (Gordon, Laumann, Gilmore, et al. 2017; Gratton et al. 2018; Seitzman et al. 2019) and may confer an advantage in task state decoding for that same person relative to other individuals who differ in their spatial network arrangement. Another possibility is that the magnitude of functional connectivity changes during tasks, rather than the spatial position, varies systematically across people.

To test whether variation in the spatial locations of networks across individuals drives differences in classifier performance, we shifted our analysis strategy. Instead of relying on common group parcellations and network definitions, we used individualized parcels and network definitions from Gordon and colleagues (Gordon, Laumann, Gilmore, et al. 2017), derived using data-driven community detection algorithms specifically for that person. We then estimated the average functional connectivity within and between networks for each state in each person, creating a set of 14×14 network correlation matrices (while differing in specific topography, 14 common networks were consistently identified across individuals - see *Methods* and **Fig. 7a**). Classifiers were trained to distinguish all tasks from rest in each participant and then tested on independent data from either that same participant or other participants. As with the other analyses reported above, individualized classifiers were able to significantly decode task from rest in both cases (Within-person test: M=.91+/−.04; Between-person test: M=.70+/− .05). Importantly, however, classifiers still performed significantly better on the same person than other people, even after matching for individualized network locations (mean effect size=.19; p<.001, **Fig. 7b**). This effect remained consistent when using group network definitions, matched for the scale of analysis (see **Fig. S10a**). This suggests that the individualized advantage for task state decoding is not driven only by differences in the spatial layout of networks, but may also be associated with differences in the magnitude of functional connectivity changes across people. We did not find an interaction effect between parcellation approach (individualize parcels compared to group parcels) and test sets (within-person compared to between person; p>.05, see **Fig. S10b**), suggesting that this effect did not strongly depend on the parcellation choice.

**Figure 7.**
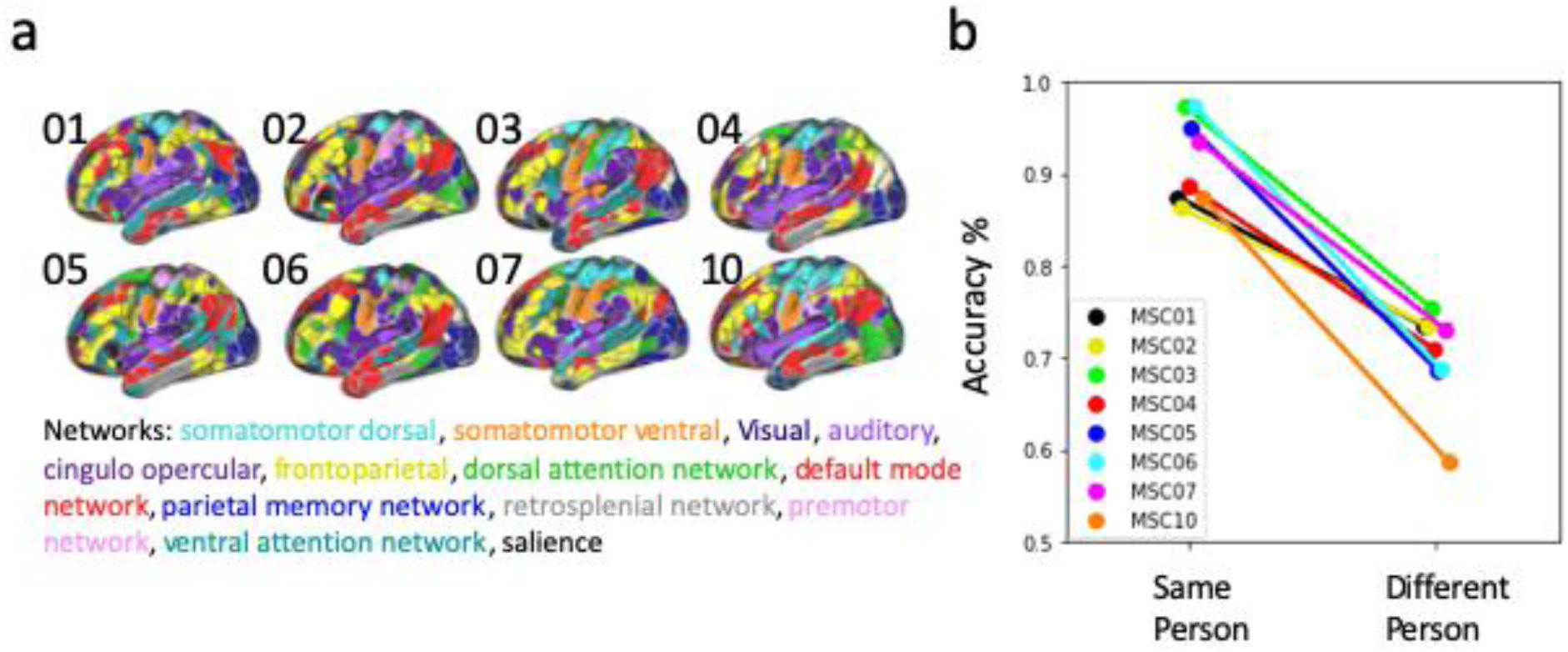
Classification performance built on individualized network parcellations. Comparison of performance of individualized classifiers on networks derived from individual parcellations. (**a**) Individual person parcellations and network definitions. FC matrices were calculated at the network level (network x network) using each participant’s network definitions in order to align across people. These FC matrices are then used to train a classifier to decode all task states from rest as in previous analyses. **(b)** Individualized classifiers based on individualized networks were significantly able to decode task from rest in both the same and new participants but performed consistently better for the same person (p<.001).

### External Replication: Within-person prediction is consistently higher

Finally, we sought to replicate our primary findings in an external, independent dataset. To this end, we analyzed a new precision dataset collected at Northwestern (data acquisition and analysis are described in the *Supplemental Methods*). This dataset currently consists of N = 22 individuals who each participated in 9 hours of MRI data collection across 4 separate sessions (along with 5+ hours of behavioral data collection outside of the scanner on other days). During each session, participants completed runs of resting-state fMRI and two other primary task conditions: a “mixed”-design task (consisting of cue-target paradigms with alternating blocks of a mental rotation, rhyme/no-rhyme judgment, and abstract/concrete auditory tasks) and a “slow-reveal” task (during which an object line drawing is slowly revealed from behind a perceptual noise mask). We estimated functional connectivity separately for each run in each task, and used these functional connectivity matrices to create individualized classifiers as in the MSC with a leave-one-session-out cross-validation procedure. As before, test sessions could be from either the same or a different participant. This procedure was repeated across each of the 22 participants. We created classifiers that distinguished between all tasks vs. rest (“All task”), each single task vs. rest, and multi-class versions that distinguished among all 3 conditions.

Performance in this new dataset (Fig. S12) was consistent with our other analyses in the Midnight Scan Club. As seen above, within-person classification performance was significantly higher than between person for the *All Task* (mean difference in accuracy = 0.35, pval < .001), *Single Task* (Mixed = 0.29, pval< .001, Slow Reveal = 0.16, p < .001), and *Multiclass* (mean difference in accuracy = 0.30, pval < .001) comparisons. This replication supports the prior findings such that differences in the ability to predict task state in within-vs between person classification are substantial. This is a notable replication as this dataset was collected on new subjects, a different scanner with fairly distinct pulse sequences, and new tasks. Thus, this replication suggests that these findings are likely fairly robust.

## Discussion

A central question in cognitive neuroscience is how brain regions coordinate their activity during complex tasks. In this study, we applied machine learning to precision fMRI data to determine the extent to which brain network changes during task states generalize across people or are individually-specific. We found that high-quality precision fMRI data led to robust individual-level classification of task state based on functional networks. These individualized classifiers were significantly above chance both in the same and in new people, indicating the presence of task generalizable effects. However, classifiers tested within the same individual had substantially higher accuracy than those tested across individuals, suggesting the presence of strong idiosyncratic features that are obscured in cross-subject analysis. This result was consistent across people, classifier procedures, and even in a new independent dataset. While single brain networks were able to predict task state alone, features for predicting task state were distributed across multiple systems; these features differed systematically across individualized classifiers and from groupwise approaches. Based on these findings, we suggest that task state is associated with systematic and idiosyncratic changes in brain networks that are often masked in cross-subject analyses. Using an individualized, high data approach can help capture these features and improve our understanding of how brain networks are altered during tasks.

### Generalizable and person-specific changes in functional networks predict task states

The brain responds in many different ways during tasks to instantiate a full ‘task state’. Here, we focus on the responses of large-scale functional networks - how activity patterns are coordinated between regions. Intrinsic functional connectivity architecture is altered in only relatively subtle ways during different task states (Cole et al. 2014; Gratton et al. 2016). However, prior work has consistently demonstrated that sensitive machine learning methods can effectively be used to predict task state from whole-brain functional connectivity data (Shirer et al. 2012; Gonzalez-Castillo et al. 2015; Madhyastha et al. 2015; Rosenberg et al. 2016; Rosenberg et al. 2018; Fong et al. 2019; Chen et al. 2020). In fact, it is possible to predict tasks designed to measure a variety of cognitive processes including attention (Alnæs et al. 2015; Gonzalez-Castillo et al. 2015; Madhyastha et al. 2015; Rosenberg et al. 2016; Rosenberg et al. 2018; Fong et al. 2019), memory (Shirer et al. 2012; Krienen et al. 2014; Braun et al. 2015; Gonzalez-Castillo et al. 2015; Jiang et al. 2020), language (Krienen et al. 2014; Jiang et al. 2020), and math processing (Gonzalez-Castillo et al. 2015) with accuracy levels well above chance. These investigations have typically been conducted using cross-subject prediction, with classifiers trained on a subset of participants and then tested on left out subjects. This approach inherently assumes that task states will induce similar network modulations across participants. Our work builds on these findings by showing that you can predict task state even using classifiers trained on a single person’s data. These individualized classifiers were able to predict task state above chance in other individuals (at similar levels to that seen with more typical cross-subject classifiers, see **Fig. S4**). These findings underscore that there are common, generalizable aspects to how brain networks are altered during tasks. However, our findings also showed a significant improvement in prediction when classifiers were used to predict state within the same person. Cross-subject classifiers can eventually reach performance levels similar to that seen with our individualized classifiers, but doing so requires ∼4x as much training data (Fig S4d). Intriguingly, even at their maximal performance, these cross-subject classifiers uncover distinct features relative to those found in our individualized classifiers (Fig. S8). This suggests that an individualized approach may help to uncover additional novel brain network features associated with task-related states that are masked in typical cross-person analyses.

In contrast to *functional connectivity* based prediction, the advantage of within-subject prediction has been recognized for a relatively long time in MVPA analyses based on fMRI *task activation* signals (Norman et al. 2006). Haxby and colleagues (Haxby et al. 2001) conducted a similar set of comparisons to the ones we included here to demonstrate the advantage of within-subject classification (relative to between-subject) for evoked representations of faces, houses and other visual category stimuli. Indeed, it has been suggested that “Because MVPA analyses focus on high-spatial-frequency (and often idiosyncratic) patterns of response, MVPA analyses are typically conducted within individual subjects.” (Norman et al. 2006). However, as noted above, many papers that conduct MVPA analyses on *functional connectivity* patterns from large-scale brain networks have focused on between-subject designs (for review see (Du et al. 2018); e.g., (Shirer et al. 2012; Alnæs et al. 2015; Madhyastha et al. 2015; Rosenberg et al. 2016; Fong et al. 2019; Jiang et al. 2020)). This may be due to a combination of historic factors such as (1) the relatively small amounts of data often collected for FC analysis (5-10 min per person), which limits the ability to do multi-run training/testing as we have provided here, and (2) that many early FC classification papers arose from the clinical literature (where the goal is typically to classify between person characteristics) rather than the cognitive literature (aiming to classify processes/representations within a person) (Dosenbach et al. 2010; Gabrieli et al. 2015; Pruett Jr et al. 2015; Siegel et al. 2016; Smyser et al. 2016; Greene et al. 2018; Steele et al. 2018; Graña and Silva 2021). It may also have been thought that (3) the scale of FC analyses (usually based on entire regions or networks) were not as subject to individual variation as the finer scale (voxel-level, within region) analyses associated with task activation MVPA. However, recent work has demonstrated pronounced individual differences even in this scale of organization, especially in association regions of the cortex (Mueller et al. 2013; Miranda-Dominguez et al. 2014; Braga and Buckner 2017; Finn et al. 2017; Gordon, Laumann, Gilmore, et al. 2017; Gratton et al. 2018; Braga et al. 2019; Seitzman et al. 2019; DiNicola et al. 2020). We hope that this work will serve as a demonstration of the importance of individual prediction models in functional network analyses, an advantage that has already been appreciated for in the prediction of evoked task patterns. An interesting avenue in future work will be to understand how these alterations in large-scale functional network architecture are related to these other characteristics of neural responses (e.g., task activations, and finer scale spatial/temporal patterns) that will, jointly, make up a neural task state.

### Sources and consequences of individual differences in task states

Here, through a number of diverse analyses, we show a consistent benefit for within-person classification in the prediction of task states. Classifiers showed significantly higher accuracy for predicting task state from data from the same person relative to new people. This finding was seen in each of the eight precision datasets we measured and was consistent across different tasks (**Fig. 2b**), networks (**Fig. 5**), subsets of samples (**Fig. S3**), versions of the classifier, (all task **Fig. 2a**, single task **Fig. 2b**, multiclass **Fig. 3a**) and classifier implementations (ridge regression, logistic regression, support vector machines, **Fig. S1**). Notably, we were able to replicate these findings using an independent precision fMRI dataset collected on a different scanner and using new tasks (Fig. S12). This performance improvement was seen even in very similar task conditions, with matched stimuli (**Fig. S5**). Consistent with past results using cross-subject approaches (Shirer et al. 2012; Krienen et al. 2014; Alnæs et al. 2015; Gonzalez-Castillo et al. 2015; Madhyastha et al. 2015; Rosenberg et al. 2016; Fong et al. 2019; Jiang et al. 2020), individualized classifiers tested on new people were able to perform significantly above chance (and similar to that seen with classic cross-subject classifiers, **Fig. S4**; see previous section), indicating that our findings were not driven by general poor performance of the individualized classifiers. Rather, the large enhancement in within-person classification suggests that individualized classifiers have the potential to reveal novel, robust features of task states that are obscured in typical cross-person designs.

These findings align with past results indicating that there are significant differences in functional brain networks across people (Mueller et al. 2013; Finn et al. 2015; Gordon et al. 2016; Braga and Buckner 2017; Gordon, Laumann, Gilmore, et al. 2017; Gordon, Laumann, Adeyemo, Gilmore, Nelson, Nico U. F. Dosenbach, and Petersen 2017; Bijsterbosch et al. 2018; Kong et al. 2019; Seitzman et al. 2019), as well as interactions between individual differences and changes in task states (Geerligs et al. 2015; Gratton et al. 2018; Xie et al. 2018). Here, we provide evidence that these individual differences have a substantive impact on task state prediction. Within-person classification may be necessary to reveal robust, impactful, but idiosyncratic features of brain network states, observations that are likely to be central to enhancing our understanding of cognitive neuroscience. The growth of precision fMRI datasets sampling a variety of task conditions such as the Natural Scenes Database (Naselaris et al. 2021), the Individual Brain Charting dataset (Pinho et al. 2018), and StudyForrest (Hanke et al. 2014; Hanke et al. 2016) will be instrumental in this regard.

What are the sources of these individual differences in classification? One possibility is that individual features may be driven by differences in the spatial layout (functional-anatomical topography) of brain networks. For many of the analyses in this manuscript, we used a group parcellation (Gordon et al. 2016). Poor between-person classification performance could therefore be driven to the extent that any individual showed deviations from this typical group architecture (Gordon, Laumann, Gilmore, et al. 2017), consistent with suggestions that spatial topography is a major form of individual differences in brain networks (Bijsterbosch et al. 2018). Deviations in spatial topography may be an important limitation in most past task state classification relying on a cross-subject prediction, but could be addressed in the future through the use of functional alignment procedures, based on matching individualized-networks (Gordon, Laumann, Gilmore, et al. 2017; Kong et al. 2019) and parcels (Glasser et al. 2013; Kong et al. 2021), or through hyperalignment, a method that has been successful at increasing model performance across participants (Haxby et al. 2011; Geerligs et al. 2015; Guntupalli et al. 2016; Guntupalli et al. 2018; Haxby et al. 2020). However, in this work, we found that a within-person advantage persisted even after functionally aligning across people based on their own intrinsic network architecture (**Fig. 7**). Thus, we interpret this result as suggesting that the within-person advantages we see in this study are more likely to be driven by idiosyncrasies in the magnitude and specific pattern of functional connectivity changes during task states, than in differences in the stable spatial layout of the networks (e.g., the extent to which frontoparietal and visual regions may be coupled during a given task, rather than where the frontoparietal network is found in a given person). Further support for this may be derived from the variable ability of the individualized classifiers to generalize to new task contexts within this dataset (Fig. S11). As the overall spatial layout of brain networks is fairly consistent across task states (Cole et al. 2014; Gratton et al. 2018; Kraus et al. 2021), one would expect that effects driven by spatial layout alone would generalize across tasks. In future work, we hope to further expand on these initial insights with additional methods that seek to match functional topography across individuals at finer scales, such as hyperalignment (Haxby et al. 2011; Guntupalli et al. 2016; Guntupalli et al. 2018; Haxby et al. 2020) and parcel-based matching methods (Kong et al. 2019).

An additional possibility is that individual task effects may be associated with differences in behavioral performance or strategy for completing tasks (Fox and Raichle 2007; Pearce and Moran 2012). As the tasks from the Midnight Scan Club were relatively simple and each participant was at ceiling performance, this hypothesis would require that even minor differences in strategies during a simple task could have measurable effects on brain networks. One exciting avenue of future work will be to evaluate whether precision data can predict diverse strategies and behavioral outcomes. Studying individualized task network features across a variety of contexts may help untangle these possibilities and piece apart the functional relevance associated with different states.

### Task state information is found across many large-scale systems

One question that arises from the current results is whether specific brain regions or connections are particularly influenced by task states. Prior work has suggested that tasks influence a range of within and between network connections, and focused on highlighting specific connections associated with various tasks (Cole et al. 2014; Gratton et al. 2016; Shine et al. 2016; Rosenberg et al. 2018). Like with past studies, we also found that both within and between network connections contributed to decoding of task state. Interestingly, however, we found that task state information was very distributed across the connectome: almost every block of network to network connections was sufficient to decode task state (within a person), but these single networks didn’t outperform random distributed feature selection when matched on feature number. These results are consistent with past work (Nielsen et al. 2020) which showed that distributed brain network features outperform features from single networks in predicting age, likely driven by the more diverse information that can be gained by sampling from many different systems. Our findings suggest that completing various tasks optimally recruits distributed systems rather than relying solely on any one network. Notably, however, the task states tested here differ in a number of ways relative to rest. Future work using more carefully matched/controlled task state comparisons may illuminate more selective patterns of modulation in functional networks. However, given the nature of functional network measures, these may still be distributed across multiple functional systems (e.g., decoding of similar task conditions in the incidental memory task still relies on features distributed throughout the brain, see Fig. S13).

Importantly, regardless of the feature selection approach (single network, block, or random feature), we found that within-person tests consistently outperformed between-person tests. This finding not only underscores the robustness of our original result but also suggests that idiosyncratic task network features are distributed throughout different brain regions. Indeed, analyses of the feature weights used in our machine learning models showed consistency of distributed network features across folds of a single person, but substantial deviations across people as well as from more standard cross-subject approaches. This suggests that individualized classifiers highlight fundamentally different features from group-level classifiers (and from each other). The next frontier will be to understand how these idiosyncratic and common features tie to specific cognitive processes.

### Applications of personalized state classifiers

Our current findings have focused on predicting task states. One interesting question for future work will be to determine whether individualized classifiers will also be useful in predicting fine-scale cognitive states related to specific aspects of memory (Avery et al. 2020), attention (Rosenberg et al. 2016), and emotion (Rohr et al. 2015), as well as broad states a person experiences such as arousal (Shine et al. 2016; Li et al. 2019) and sleep (Tagliazucchi and Laufs 2014; Tagliazucchi and van Someren 2017). If it is broadly true that idiosyncratic features are important, then person-specific precision analyses such as the ones conducted in this study may uncover new important features across a range of domains. As an initial hint in this direction, we found in this work that individualized classifiers are able to distinguish among even very similar conditions within a single task (e.g., repeated presentations of stimuli within the memory task, see Fig. S5). Notably, these conditions only differ in their repeated nature, with equal perceptual and explicit task demands. Based on these observations, we hypothesize that our findings could be applied to more finely-subdivided cognitive states.

Given the highly replicated finding of individual variability in functional network topography across individuals (Finn et al. 2015; Gordon et al. 2016; Gordon, Laumann, Gilmore, et al. 2017; Gordon, Laumann, Adeyemo, Gilmore, Nelson, Nico U. F. Dosenbach, and Petersen 2017; Braga et al. 2019; Seitzman et al. 2019), including in large community-based samples (Mueller et al. 2013; Bijsterbosch et al. 2018; Kong et al. 2019; Seitzman et al. 2019), it is likely that individualized classifiers may have unique advantages across a variety of contexts. Clinical applications may ultimately require a balance between precision applications which may offer unique advantages in identifying idiosyncratic features of diverse states and classifiers built to maximize cross-person generalization (e.g., large datasets with less precise measurements which can be used to capture large-scale generalizable features across a population (Satterthwaite et al. 2014; Marek et al. 2020)).

One exciting avenue for future research will be to understand whether there are subsets of individuals that share idiosyncratic features (Seitzman et al. 2019). An understanding of the properties of joint (or disjoint) variation across subsets of people might then be used to improve cross-subject decoding while preserving sensitivity to individual differences. Uniting individualized approaches and targeted cross-subject studies may help to maximize our knowledge and prediction power.

### Limitations and practical considerations for individualized task state prediction

By necessity, individualized classifiers were built on a relatively small number of samples (80) and sessions per participant (10). However, the presence of eight participants allowed for eight independent models to replicate the individualized classifier results. We were further able to replicate these findings in N = 22 precision fMRI datasets collected an independent dataset collected on a different scanner and using different tasks. These individualized classifiers showed many consistent results, including significant ability to decode task state within and across people, across tasks, and across feature subsets. Prior work has shed light on the effects of data quantity and reliability in FC matrices (Laumann et al. 2015; Noble et al. 2017). To study how data quantity affects individualized classifiers, we iteratively increased the number of samples (16-80 task/rest pairs) used in our analysis. We found that increasing the number of samples in the training set influenced the models ability to accurately decode tasks from rest within a person, plateauing around 25 sample pairs (50 samples). However, the number of samples only modestly improved the model’s ability to predict state in a new person. We found that increasing the number of training sets for cross-subject classifiers also improved model performance (Fig. S4). However, nearly 4 times as much training data was needed to achieve comparable performance to the individualized classifiers. Due to the nature of our sample, this was tested by increasing the number of training samples from each participant (while keeping the number of participants constant), suggesting that precision fMRI datasets may also lead to improvements in cross-subject decoding. Future work will be needed to compare within and between training sample contributions to cross-subject decoders.

Like with sample increases, we also found that increasing numbers of features improved classification, even when drawn randomly from different positions in the functional connectome. This was particularly true for within-person classification, which showed substantial performance improvements plateauing at ∼1000 features. This finding suggests that caution should be warranted in interpreting differences in classification between networks of different sizes. Notably, these results provide guidelines for the practical application of building individualized classifiers in future studies seeking to maximize within and between person prediction.

## Conclusion

In this study, we asked whether people differ in how their brain interactions are altered by task state. Using a machine learning approach, we find that it is possible to classify task state from a classifier trained on a single individual’s multi-session data. Classification was successful both for independent data from that person as well as for new people, but a big boost in accuracy was seen for classification within-person. This result was consistent across datasets, tasks, and classifier approaches. Feature selection analyses demonstrated that task state information is distributed across large-scale systems, with single networks performing well at classifying task state but not out-performing randomly selected features. These findings have important implications for understanding how task state relates to brain networks, emphasizing the importance of considering individual differences to reveal the full magnitude of variations that are present.

## Acknowledgements

This work was supported by NIH grants R01MH118370 (CG) and 2T32MH067564 (AP) and NSF CAREER 2048066 (CG). Special thanks to Babatunde Adeyemo, Steven E. Petersen, and members of the Gratton, Dosenbach, and Greene Labs for feedback on various stages of this project.

