## Supplement for "Masked features of task states found in individual brain networks"

#### Supplemental materials

### Supplementary Methods

#### *Replication dataset: iNetworks*

##### *Dataset overview*

The iNetworks dataset includes data from 22 individuals (15 females, ages 21-33), each with four fMRI sessions completed within 6 weeks (SD=3.8) for all participants. In each session, fMRI data included resting-state scans and four tasks (see Task Designs and Analysis). Participants provided informed consent and procedures were approved by Northwestern University Institutional Review Board. All participants who had completed data collection, quality control, and processing at the time of writing this manuscript were included in the analyses.

##### *MRI Acquisition*

MRI data were acquired on a 3T Siemens Prisma at Northwestern University using a 64-channel head coil. Two T1-weighted images (sagittal, 208 slices, 0.8 mm isotropic resolution, TE1 = 1.86 ms, TE2 = 3.78 ms, TR = 2.3, TI = 1.18s, flip angle = 7 degrees) and one T2-weighted image (sagittal, 208 slices, 0.8 mm isotropic resolution, TE = 566 ms, TR = 3.2s) were collected for each participant across three separate days. Functional MRI data was collected using a gradient-echo EPI BOLD sequence (TE = 25ms, TR = 1.1 s, flip angle = 80, voxels = isotropic 2.5mm$^3$, 60 axial slices, multiband acceleration factor = 4) during each of 4 sessions. The same sequence was used for task and resting-state data. Using the same parameters, a gradient echo field map was acquired during each session for de-warping of the functional data.

##### *Functional MRI Preprocessing*

Results included in this manuscript come from preprocessing performed using fMRIPrep 20.2.0 (Esteban et al. 2018; Esteban et al. 2019; RRID:SCR_016216), which is based on Nipype 1.5.1 (Gorgolewski et al. 2011; Esteban et al. 2018; RRID:SCR_002502). A total of 2 T1-weighted (T1w) images were found within the input BIDS dataset. All of them were corrected for intensity non-uniformity (INU) with N4BiasFieldCorrection (Tustison et al. 2010), distributed with ANTs 2.3.3 (Avants et al. 2008, RRID:SCR_004757). The T1w-reference was then skull-stripped with a Nipype implementation of the antsBrainExtraction.sh workflow (from ANTs), using OASIS30ANTs as target template. Brain tissue segmentation of cerebrospinal fluid (CSF), white-matter (WM) and gray-matter (GM) was performed on the brain-extracted T1w using fast (FSL 5.0.9, RRID:SCR_002823, (Zhang et al. 2001)). A T1w-reference map was computed after registration of 2 T1w images (after INU-correction) using mri_robust_template (FreeSurfer 6.0.1, (Reuter et al. 2010)). Brain surfaces were reconstructed using recon-all (FreeSurfer 6.0.1, RRID:SCR_001847, (Dale et al. 1999)) and the brain mask estimated previously was refined with a custom variation of the method to reconcile ANTs-derived and FreeSurfer-derived segmentations of the cortical gray-matter of Mindboggle (RRID:SCR_002438, (Klein et al. 2017). Volume-based spatial normalization to a standard space (MNI152NLin6Asym) was performed through nonlinear registration with antsRegistration (ANTs 2.3.3), using brain-extracted versions of both T1w reference and the T1w template. The following template was selected for spatial normalization: FSL's MNI ICBM 152 non-linear 6th Generation Asymmetric

Average Brain Stereotaxic Registration Model ((Evans et al. 2012), RRID:SCR_002823; TemplateFlow ID: MNI152NLin6Asym).

For each of the BOLD runs found per subject (across all tasks and sessions), the following preprocessing was performed. First, a reference volume and its skull-stripped version were generated using a custom methodology of fMRIPrep. A B0-nonuniformity map (or fieldmap) was estimated based on a phase-difference map calculated with a dual-echo GRE (gradient-recall echo) sequence, processed with a custom workflow of SDCFlows inspired by the epidewarp.fsl script and further improvements in HCP Pipelines (Glasser et al. 2013). The fieldmap was then co-registered to the target EPI (echo-planar imaging) reference run and converted to a displacements field map (amenable to registration tools such as ANTs) with FSL's fugue and other SDCflows tools. Based on the estimated susceptibility distortion, a corrected EPI (echo-planar imaging) reference was calculated for a more accurate co-registration with the anatomical reference. The BOLD reference was then co-registered to the T1w reference using bbregister (FreeSurfer) which implements boundary-based registration (Greve and Fischl 2009). Co-registration was configured with six degrees of freedom. Head-motion parameters with respect to the BOLD reference (transformation matrices, and six corresponding rotation and translation parameters) are estimated before any spatiotemporal filtering using mcflirt (FSL 5.0.9, (Jenkinson et al. 2002). The BOLD time-series (including slice-timing correction when applied) were resampled onto their original, native space by applying a single, composite transform to correct for head-motion and susceptibility distortions. These resampled BOLD time-series will be referred to as preprocessed BOLD in original space, or just preprocessed BOLD. The BOLD time-series were resampled into standard space, generating a preprocessed BOLD run in MNI152NLin6Asym space. First, a reference volume and its skull-stripped version were generated using a custom methodology of fMRIPrep. All resamplings can be performed with a single interpolation step by composing all the pertinent transformations (i.e. head-motion transform matrices, susceptibility distortion correction when available, and co-registrations to anatomical and output spaces). Gridded (volumetric) resamplings were performed using antsApplyTransforms (ANTs), configured with Lanczos interpolation to minimize the smoothing effects of other kernels (Lanczos 1964). Non-gridded (surface) resamplings were performed using mri_vol2surf (FreeSurfer). As in the MSC, the native freesurfer surface was registered to fs_LR_32k space following a procedure based on (Glasser et al. 2013).

Functional data was then denoised, mapped to the surface, and smoothed using the same pipeline described in *Functional Connectivity Pre-processing* (Methods; note that in this case, the filtered FD approach was used for all participants, with a criteria of fFD < 0.1 Hz. applied for censoring (Fair et al. 2020)). The Gordon parcellation (Gordon et al. 2016) was used to define brain regions and functional connectivity was estimated using timeseries correlations among the denoised, censored timeseries. For task data, task GLM models (see next section) were applied to the data prior to the denoising step to remove evoked signals from the timeseries as in the MSC; residuals of these models then underwent the same steps of processing.

***Task Designs and Analysis***

Functional MRI data were collected during five conditions described briefly below. Task activations were modeled with a generalized linear model (GLM) using AFNI (Cox 1996). As with the MSC dataset, GLM residuals were used for time-series correlations. Task blocks/runs of the same condition were concatenated together for each session prior to creation of FC matrices.

Resting-state: Each MRI session included several resting-state scans, wherein participants were asked to stare at a fixation on the center of a black screen. Rest runs lasted approximately 5 minutes (270 TRs) each, interleaved with task runs in each session. For each subject, an average of 155.6 minutes (SD=6.3) of resting-state data was collected in total across sessions.

Rhyme Task: Each session included 3-4 runs (33 min) of a mixed block/event-related design using cue/task paradigms modeled after tasks described by (Neta et al. 2014; Gratton et al. 2016). In each run there were three blocks per run, one for 'rhyme', one for abstract/concrete (see below in Abstract/Concrete), and one for mental rotation (see below in Mental Rotation). Task blocks began with a 1.1s cue indicating which task was to be conducted in the following block; after this 18 individual trials were presented. The rhyme task (Neta et al. 2014) was a rhyming discrimination task wherein participants were visually presented with two words and asked to identify if they did or did not rhyme. There were 8 rhyme, 8 no rhyme, and 2 ambiguous trials per block. Trials consisted of words presented for 2.2s with jittered 2.2, 4.4, or 6.6 intervals (mean intertrial interval [ITI] 4.4 s). After, participants were presented with a cue (1.1s) indicating the end of a block, with 39.6s fixation periods separating each block.

The rhyme, abstract/concrete, and mental rotation tasks (see below for details on other tasks) were modeled together in a single mixed block/event-related GLM. Separate regressors were included for each task block (sustained activation) and for events (start and end cues in each task, correct and incorrect trials of different types). Event-related trials were modeled using AFNI's 3dDeconvolve with tent basis functions, producing a 15-timepoint response estimation for each condition. Entire task blocks were modeled with AFNI's 3dDeconvolve function under a 104.4-second square wave. GLM residuals used for time series correlations were produced by adding the residual error timeseries to the baseline constant for all voxels.

Abstract/Concrete Task: Abstract/Concrete task blocks were interleaved with the rhyme and mental rotation tasks in the same mixed runs and followed the same timing structure and analysis as the rhyme task. In the abstract/concrete task (Neta et al. 2014), individual trials consisted of a noun presented audibly through headphones, where participants were asked to identify if the word was an abstract or concrete noun. There were 8 abstract, 8 concrete, 2 ambiguous trials per block.

Mental Rotation Task: Mental rotation task blocks were interleaved with the rhyme and abstract/concrete tasks in the same mixed runs and followed the same timing structure and analysis. For the mental rotation task (Dubis et al. 2016; Gratton et al. 2016), white, 2D Tetris-like shapes made of 7 squares were presented on a black screen. Two stimuli were presented on either side of a central, white fixation cross. Participants determined whether the 2 stimuli, rotated with respect to each other, had the same or mirror orientation (50% probability). Eight different stimulus shapes were used. Orientation bins of 40°–60°, 100°–120°, and 150°–170° were used. To shorten reaction times, 1 of the 2 stimuli was always upright (0° rotation).

Slow Reveal Task: Participants completed a total of 3-4 runs per session (24 or 32 min.) of a perceptual recognition task using picture stimuli (based on (Ploran et al. 2007; Gratton et al. 2017)), with 21 trials per run. In each trial, stimulus revelation occurred over eight discrete steps, each corresponding with acquisition of a whole-brain image. The revelation steps occurred every 2.2 s with a between-trial jitter of 2.2, 4.4, or 6.6 s (mean intertrial interval [ITI] 4 s). At trial onset, pictures were covered by a black mask. The mask partially dissolved at each successive 2.2 s interval (i.e., revelation step) until pictures were completely revealed. Participants were instructed to press a button when they could identify the picture with a reasonable degree of confidence. When stimuli were fully revealed, participants pressed the same button again only if their earlier recognition had been correct ("Verification of Accuracy"). The slow reveal task activations were modeled with a GLM using AFNI's 3dDeconvolve function, with 7 regressors: errors of commission, errors of omission, and correct trials binned into five conditions (correct response on steps 1-3, correct on step 4, correct on step 5, correct on step 6, correct on step 7). The tent basis function was used to produce a 27-time point response estimation for trials of each condition. GLM residuals used for time series correlations were produced by adding the residual error timeseries to the baseline constant for all voxels.

#### Supplemental Tables

| Training Task | Same Person | | | Different Person | | | Within vs. between significance |
|---|---|---|---|---|---|---|---|
| | Accuracy | TPV | RPV | Accuracy | TPV | RPV | |
| All Tasks vs. Rest | .98 (.001)*** | .97(.001) | .99(.0008) | .73(.01)*** | .69(.01) | .93(.01) | p<.001 |
| Individual Networks, All Tasks vs. Rest | .91(.005)*** | .86(.008) | .88(.008) | .70(.01)*** | .76(.02) | .71(.01) | p<.001 |
| Multiclass | .83(.01)*** | | | .53(.02)*** | | | p<.001 |
| Coherence vs. Rest | .87(.02)*** | 1(0) | .81(.02) | .58(.01)*** | .98(.009) | .55(.01) | p<.001 |
| Memory vs. Rest | .98(.009)*** | 1(0) | .97(.01) | .88(.01)*** | .89(.01) | .93(.01) | p<.001 |
| Motor vs. Rest | .98(.009)*** | 1(0) | .96(.01) | .82(.01)*** | .99(.003) | .78(.02) | p<.001 |
| Semantic vs. Rest | .94(.01)*** | 1(0) | .90(.01) | .66(.01)*** | .99(.001) | .62(.01) | p<.001 |

**Table S1**. Summary table of classification performance (accuracy=M(SE)) on cross-person tests, with training on one person and testing on either the same person or a different person. Task predictive value (TPV) represents the classifier's ability to accurately label task divided by any labeling of task. Rest predictive value (RPV) represents the classifier's ability to accurately label rest divided by any labeling of rest. Model significance was determined relative to a random null created through permutation testing *p<.05, **p<.01, ***p<.001. The final column shows the difference in model performance for within-person and between-person tests (computed with permutation testing)). All Task vs. rest: binary classifier trained to distinguish all tasks from rest. Individual Network All Tasks vs Rest: similar to all task vs. rest, but used networks derived from individual parcellations (Fig. 6). Multiclass: classification models trained to distinguish between all five states (coherence, memory, motor, semantic, and rest). Coherence, memory, motor, semantic: classifiers built to distinguish a single task from rest. These analyses are described in detail in the Methods.

#### Supplemental Figures

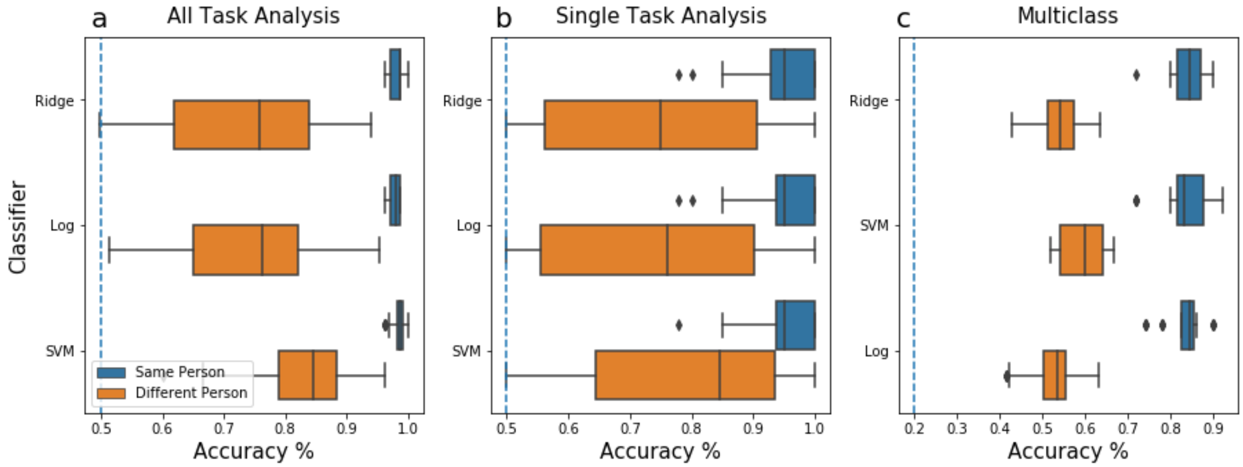


**Figure S1. Model performance across different algorithms.** Task state prediction performance across different machine learning algorithms when tested on independent data from the same (blue) or a different person (orange), shown as box and whisker plots. The whiskers represent the upper and lower quartile to the highest/lowest value that is within the 25th and 75th percentile. Points outside the range (solid diamonds) reflect outliers. Three different classifiers were tested: ridge regression ("ridge"; the primary method in the main text), logistic regression ("log"; with L2 regularization and lbfgs solvers) and support vector classification ("SVM"; using LinearSVC with default parameter settings). **(a)** Classifier performance for data tested on the same or different person when trained to discriminate between all tasks and rest data for a person. **(b)** Classifier performance for classifiers trained to distinguish a single task from rest (averaged across tasks). **(c)** Multiclass performance for classifiers trained to distinguish among all four tasks and rest. Performance was similar across algorithms, with a consistent benefit for within-person tests relative to between-person tests.

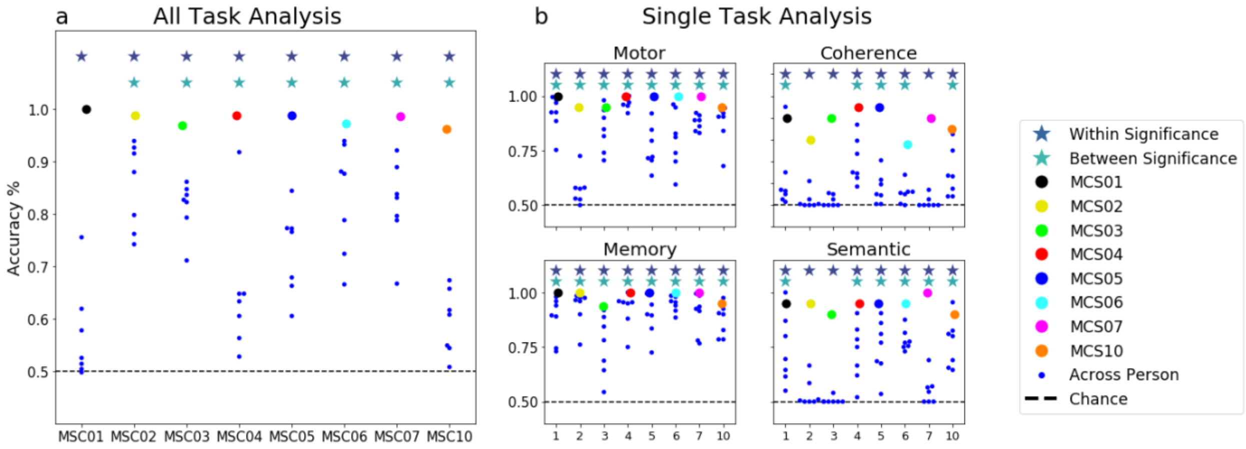


**Figure S2. Model performance per person.** Comparison of model performance for models trained on a particular individual (x-axis) and tested on either the same individual (large colored dots) or other individuals (blue dots). Figures depict the average accuracy across folds, with chance (50%) represented as a dashed black line. Significance per individualized classifier plotted as stars (dark blue within, light blue between; based on comparing true scores to a permuted null; $*p<0.05$). (**a**) Task state classification performance for classifiers trained to distinguish between all tasks and rest **(b)** Classification performance for classifiers trained to distinguish single tasks from rest. In all cases, performance is significantly higher when testing on the same person compared to between people.

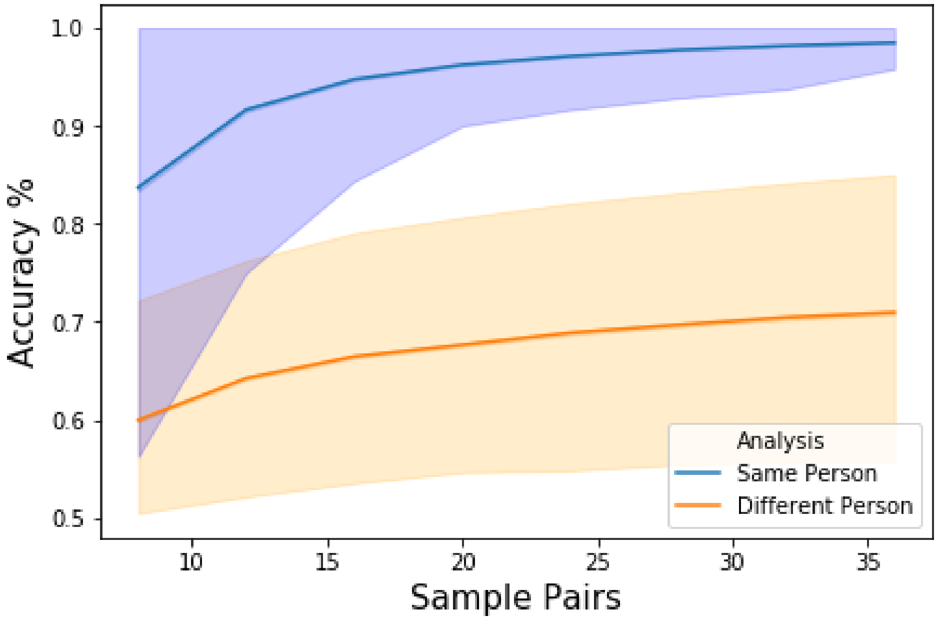


**Figure S3**. **Data Quantity Analysis**. Classifier performance on discriminating all tasks vs. rest for data tested on the same (blue) or a different person (orange). Training samples were increased iteratively in pairs (rest and task) from 16 samples to 80 samples. This process was repeated with 1,000 random training sample iterations. Error bars represent the 5th and 95th percentile at each sample pair. In all cases, classifier accuracy increased with larger numbers of samples. Greater improvements were seen when testing on the same person compared to testing on a different person.

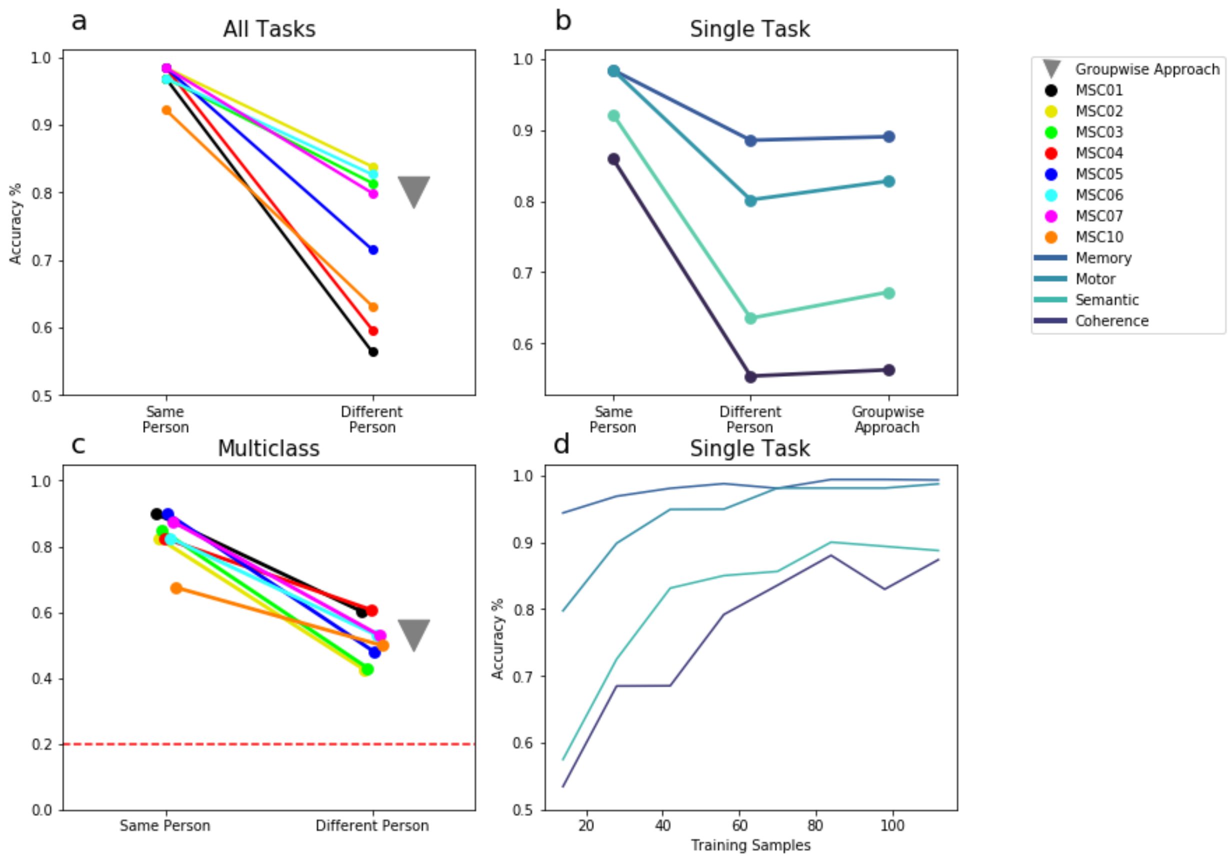


**Figure S4. Comparison of individualized models to standard "groupwise" classification. (a)** Average accuracy of classifier performance in discriminating all tasks from rest using individualized classifiers tested on the same (left) or new (right) people (as in the main text) or a more standard "groupwise" approach (gray triangle). The groupwise classifier was trained using a leave-one-subject-out cross-validation procedure and tested on the left-out subject (see Methods). All versions were trained and tested on the same number of exemplars (in this case, 7 matrices in each training set, 1 matrix in each test set for the Single Task and Multi Class Analyses, 28 matrices in each training set, 4 matrices in each test set for the All Task Analysis). **(b)** Similar to A, but for classifiers built to distinguish a single task from rest **(c)** Similar to A, but for multi-class classifiers built to discriminate among all tasks and rest. In each of these cases, group-wise classification (gray triangles) was similar to the classification seen for individualized classifiers tested on other people, below the level seen for individualized classifiers tested on the same person. **(d)** We next asked whether adding additional training data could improve group-wise classification performance to similar levels as seen within individualized classifiers. Plots show average classifier performance for distinguishing all tasks from rest using the groupwise approach when incrementally including more training data (given the nature of our dataset, this additional data represents more sessions of data from each participant rather than new participants). Groupwise classifiers reach similar performance as individualized classifiers when over 70 samples are used in training, approximately 4x as much data as is used in the individualized classifier analyses.

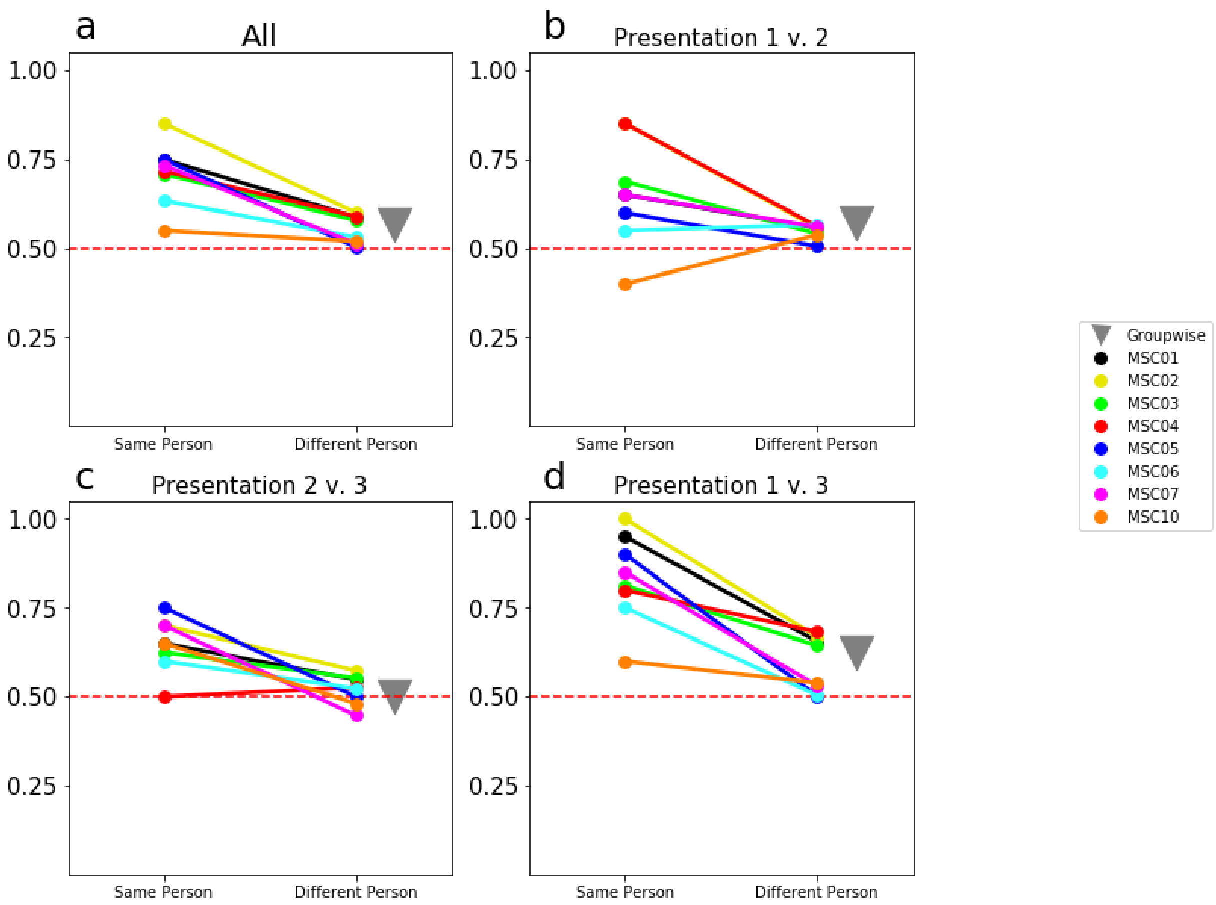


**Figure S5. Binary classification across similar task conditions.** Performance of individualized functional connectivity classifiers on classifying among similar task conditions. Here we present average accuracy when training the classifiers to discriminate among similar conditions from the memory task, specifically contrasting blocks with the first, second, or third repetition of presented stimuli. **(a)** Average performance of individualized classifiers at distinguishing among memory conditions when tested on the same or different people. **(b)** Classifier performance on distinguishing between presentation 1 v. presentation 2, **(c)** presentation 2 v. presentation 3 or **(d)** presentation 1 v. presentation 3. For comparison, we also included a groupwise approach for each analysis, in which an equal amount of data sampled across different subjects was used for training (gray triangle). Notably, in all conditions participants were viewing the same stimuli and performing the same explicit task (a categorization judgment); the only difference between conditions was the number of previous repetitions of the stimuli (Gordon, Laumann, Gilmore, et al. 2017). All functional connectivity classifiers were able to discriminate these conditions above chance (p<.001), but as in the other analyses from the manuscript, the within-participant classification significantly outperformed between participant classification (p<.001) even for these task conditions with high similarity.

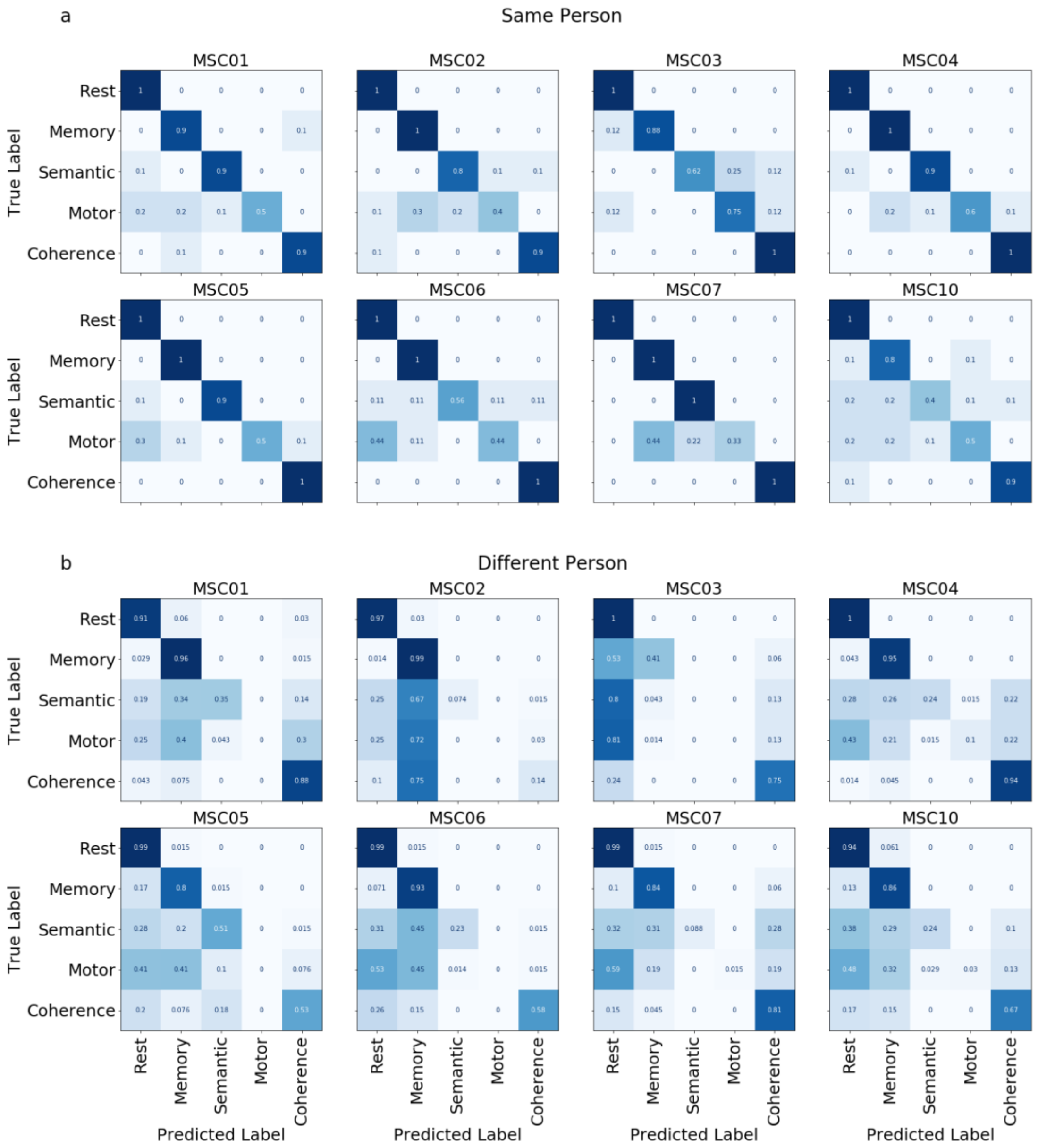


**Figure S6. Multiclass confusion matrices of individualized classifiers.** Multiclass output of confusion matrices for each individualized classifier when testing on left out sessions from either **(a)** the same person or **(b)** a different person. Performance was relatively high and similar across individualized classifiers. In all cases, within-subject tests showed a strong diagonal (correct performance) with a low number of errors. Between-person, tests were more varied, with biased errors in classification (e.g., for motor and semantic conditions especially).

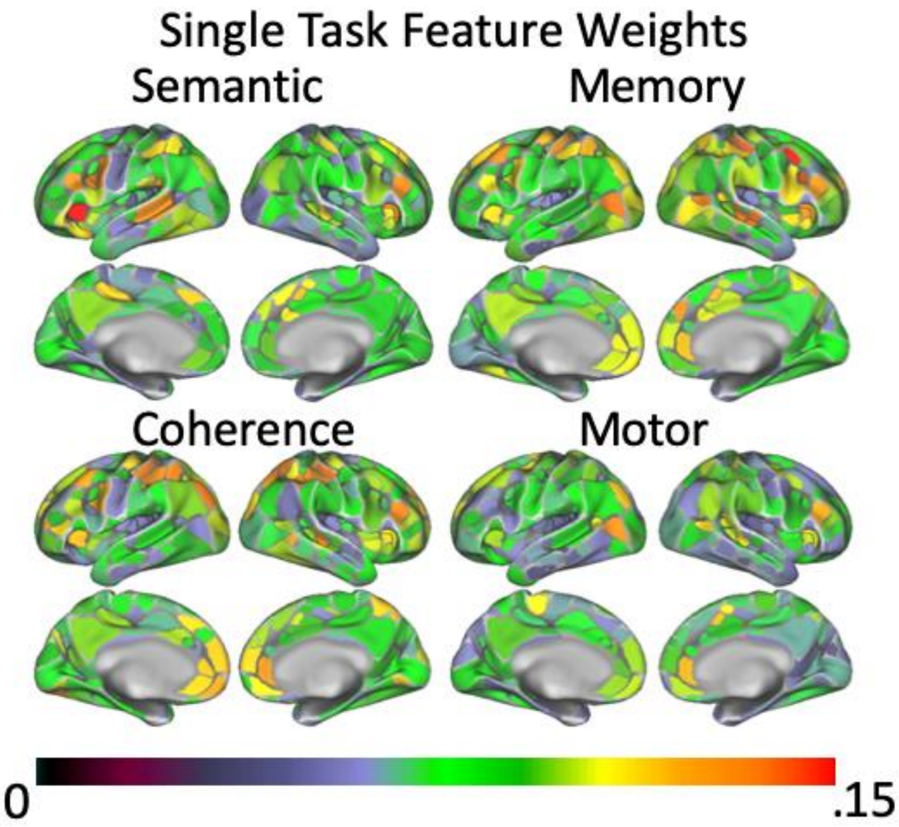


**Figure S7. Feature Weights Across Task.** Absolute average feature weights on the single task analysis for a single person (MSC05), as in Fig. 4. Feature weights vary by task.

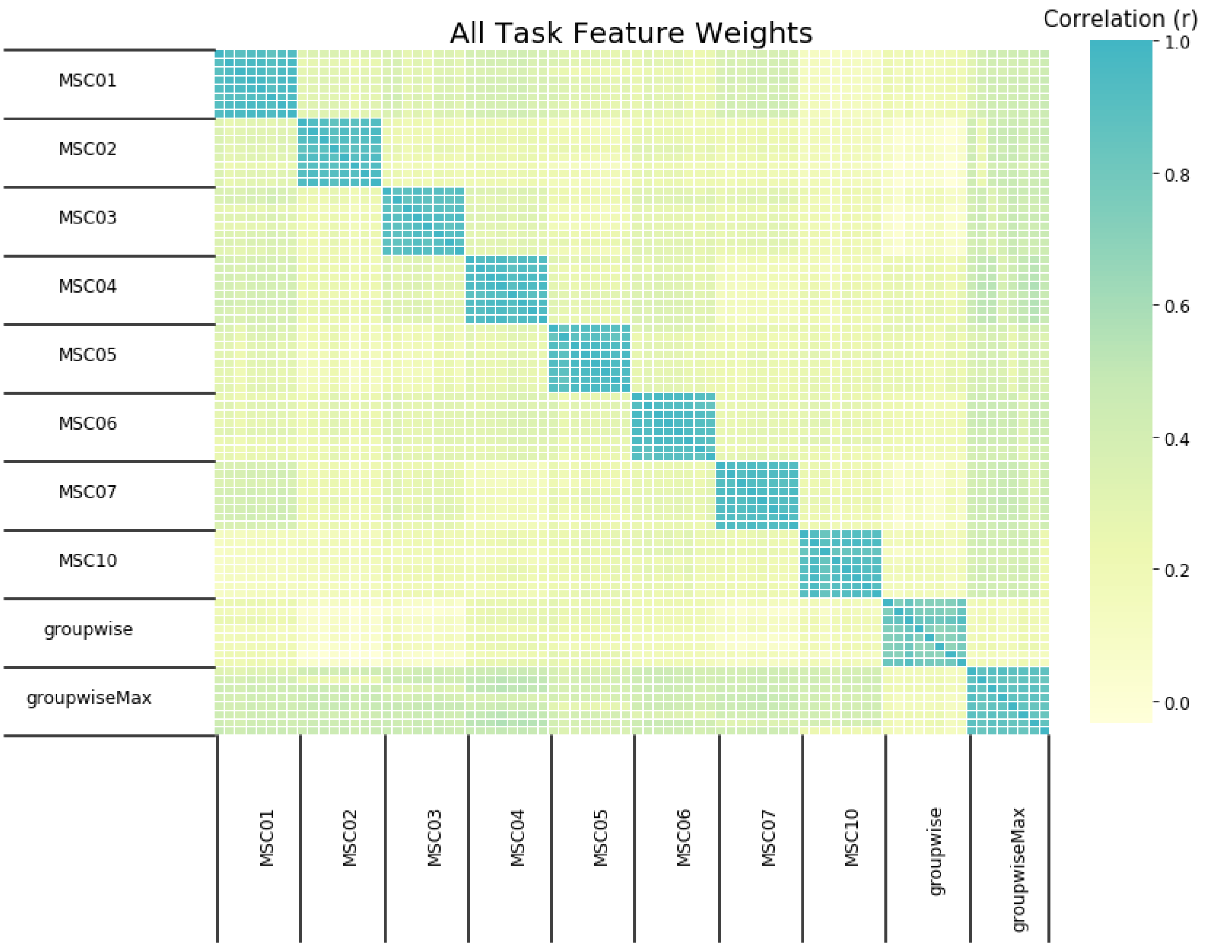


**Figure S8. Comparison of Feature Weights.** Here we took the correlation of feature weights from each fold of classification for both individualized (training data from a single person) and group-wise classifiers (training data from many subjects). We included both a "groupwise" classifier with matched amounts of training data to our individualized classifiers and a "groupwiseMax" classifier with all possible data across people (7x as much training data as the others). Analyses were based on the All Task vs. Rest (Binary) analysis. Feature weights were much more highly correlated across folds within a person than across people or to the groupwise approaches

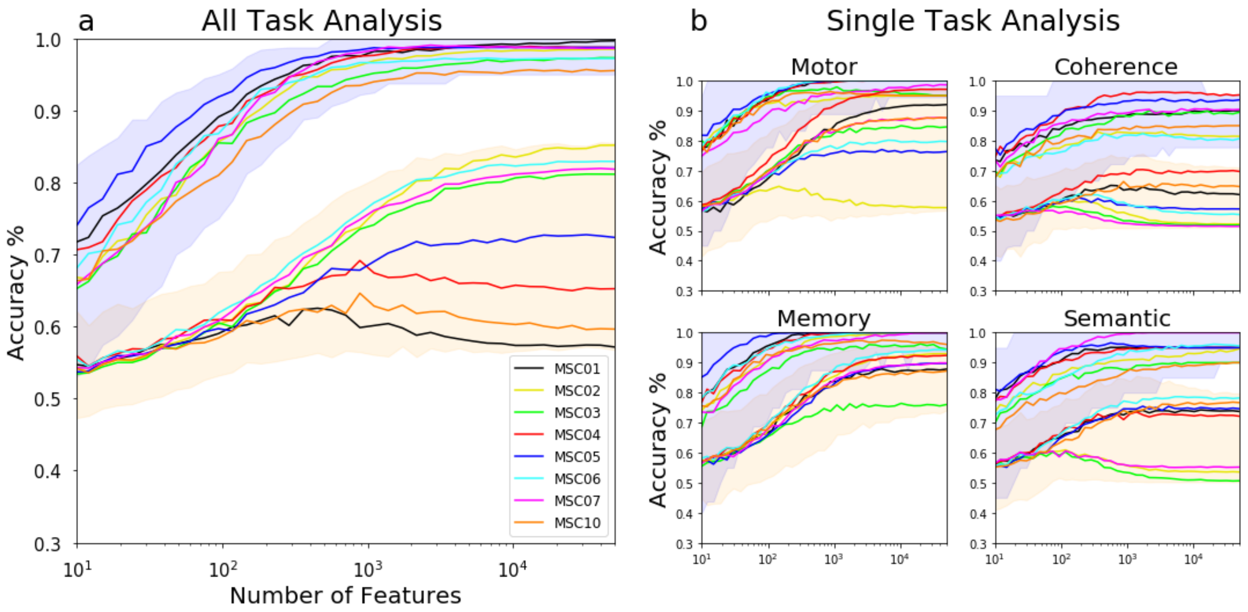


**Figure S9. Random feature selection split up by individuals.** Here we present random feature selection (10 - 50,000 number of features) when training the classifiers to (a) discriminate all tasks from rest or (b) single tasks from rest. We've plotted how individual test sets perform when tested on the same person as the training set (95% range in blue) compared to testing along a different person (orange). The mean for each individual is shown in a colored line within each set.

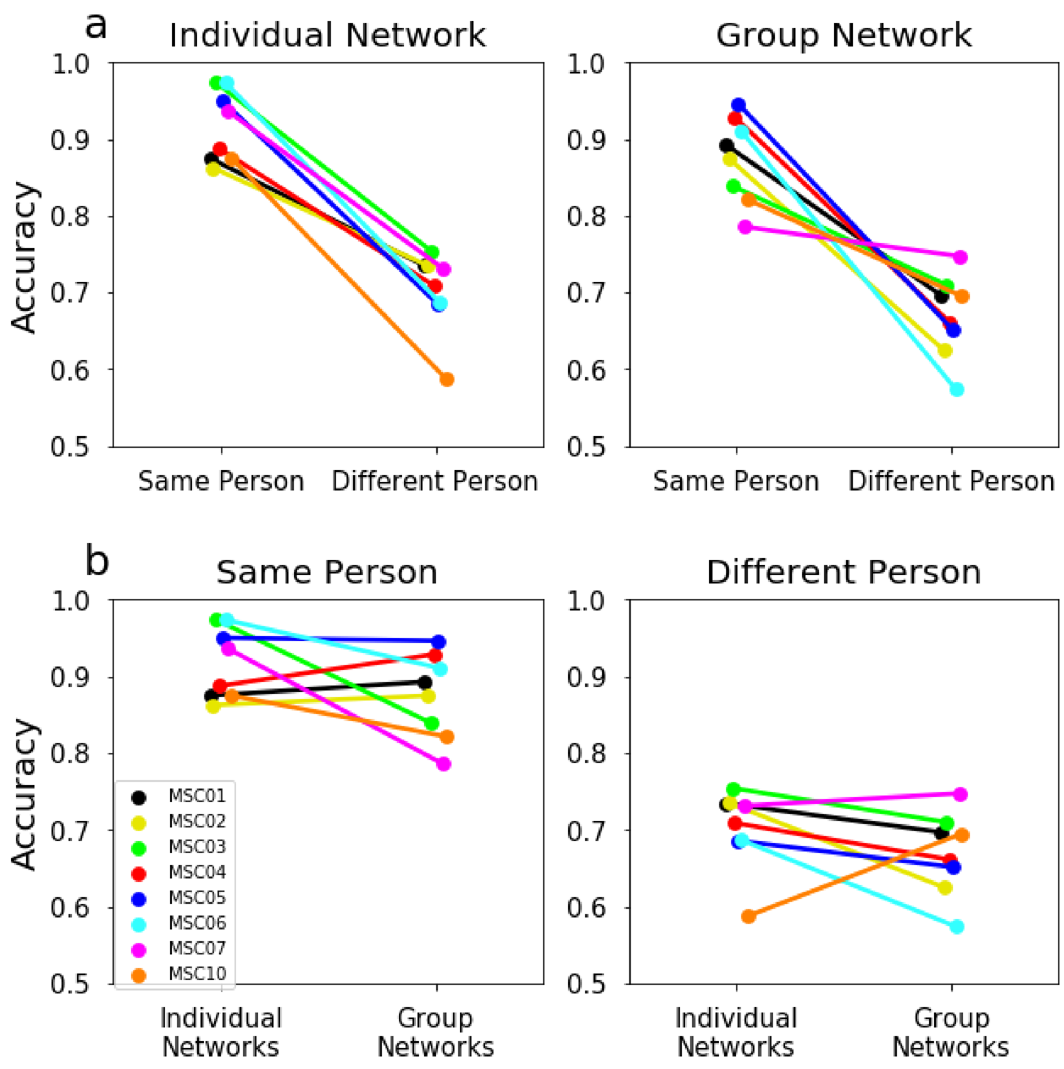


**Figure S10. Network level classification.** Classification performance for discriminating tasks from rest, analyzed at the network level for networks generated from **(a)** (**left**) individualized parcellations (functionally matched across participants) and (**right)** group network parcellations. In each case, performance is higher within-person compared to between-person, suggesting that the within-person advantage is not driven by stable differences in the layouts of brain networks. (b) Additionally we compared parcellation approach for each test set separately and did not find a significant interaction effect between parcellation approach and test sets.

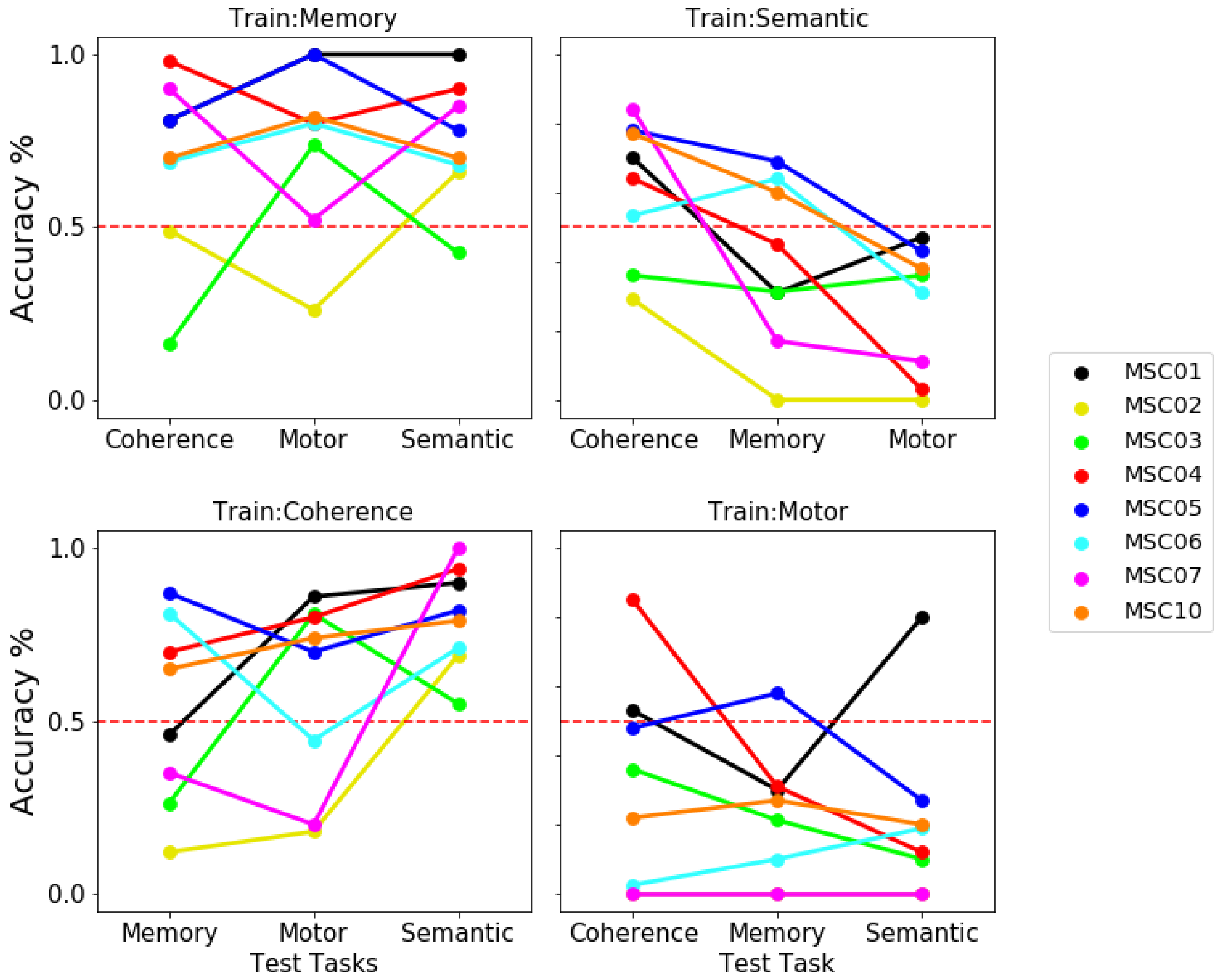


**Figure S11. Between Task Classification.** Here we tested individualized classifiers trained on a single task comparison (e.g., memory vs. rest) on unseen tasks (e.g., motor, semantic, coherence) from the same person. Classifiers built on a single task vs. rest sometimes show the ability to generalize to new tasks, but can also show large biases in their classification, and this is variable across task pairings and subjects. These biases may be due to overlapping constructs/demands across the tasks (e.g., all of the tasks have visual input and motor responses), but the tasks also differ from one another in other ways that are less psychologically interesting (e.g., he memory task was the longest, the motor task lost the most frames due to task-related movement confounds, the semantic and coherence tasks were collected in close temporal proximity to one another). Future work will be needed to better understand how individualized task classifiers generalize across new conditions.

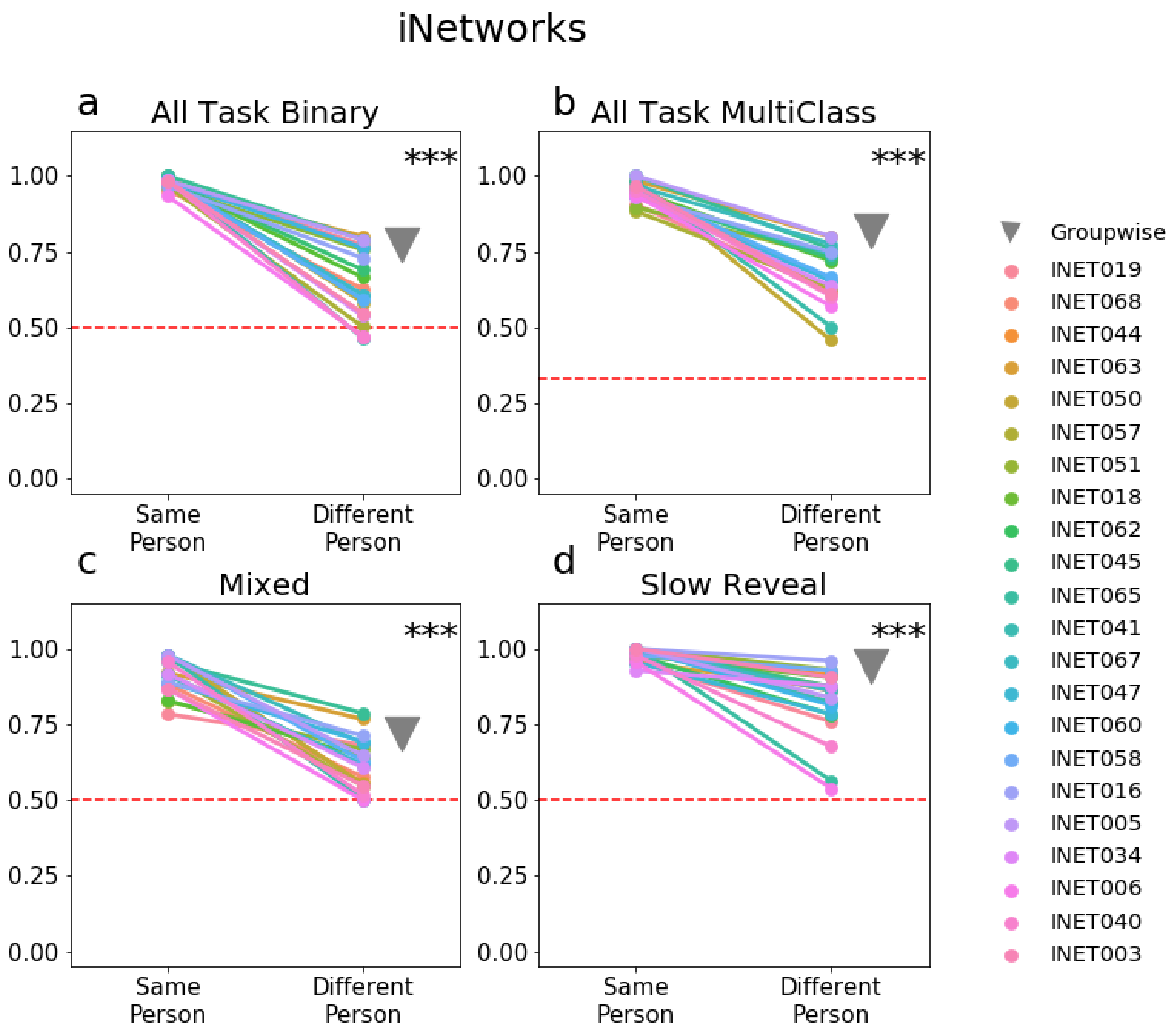


**Figure S12. Replication of individualized classification from an independent dataset.** We replicated our results from the MSC in a new dataset ("iNetworks") collected at Northwestern University in N = 22 participants, who each completed 9 hours of MRI with runs of resting-state, a "mixed" set of cue-target paradigms, and a decision-making "slow-reveal" task (see Supplemental Methods for task details). As in the main text, a subset of each individual's data was used to train an individualized classifier to distinguish task states. This classifier was then tested on independent data from that same participant or other participants. As with the MSC, we found that individualized classifiers were successful in decoding task state across a range of conditions, but that performance was significantly higher for the same person relative to new participants (*** $p < 0.001$ for within vs. between classification, gray triangle = groupwise leave-one-subject-out classification).

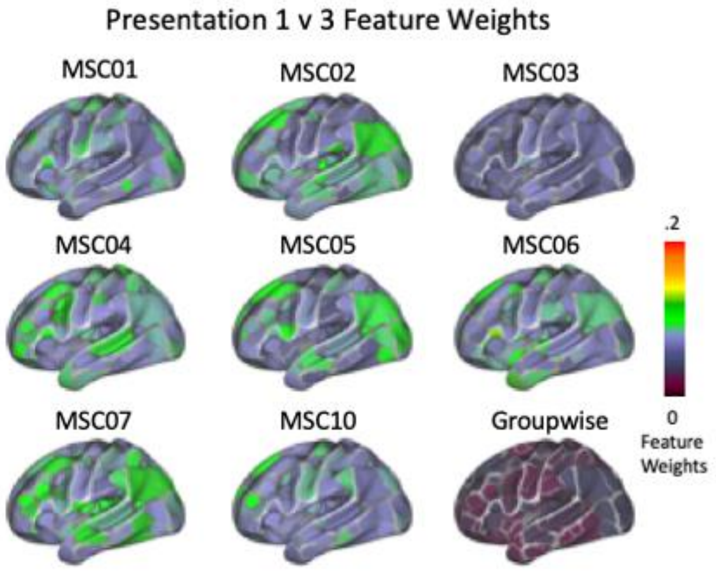


**Figure S13. Average feature weights for classification of repeated presentations in the memory task.** We present average feature weights for each individualized classifier compared to the groupwise approach when training classifiers to discriminate among similar conditions from the memory task, specifically contrasting blocks with the first and third repetition of presented stimuli. Feature weights were overall lower and appeared to involve fewer regions, as might be expected for this more selective contrast. However, consistent with results of the broader task comparisons, widespread regions across cortex had strong feature weights for this analysis. Weights were also distinct across individuals, and generally higher within a person than in the groupwise approach. Thus, this more specific analysis supports the idea that individualized classifiers identify a range of broadly distributed and individually-specific features that differ from class groupwise approaches, even with more tightly controlled comparisons.
